# The p97/VCP adapter UBXD1 drives AAA+ remodeling and ring opening through multi-domain tethered interactions

**DOI:** 10.1101/2023.05.15.540864

**Authors:** Julian R. Braxton, Chad R. Altobelli, Maxwell R. Tucker, Eric Tse, Aye C. Thwin, Michelle R. Arkin, Daniel R. Southworth

## Abstract

p97/VCP is an essential cytosolic AAA+ ATPase hexamer that extracts and unfolds substrate polypeptides during protein homeostasis and degradation. Distinct sets of p97 adapters guide cellular functions but their roles in direct control of the hexamer are unclear. The UBXD1 adapter localizes with p97 in critical mitochondria and lysosome clearance pathways and contains multiple p97-interacting domains. We identify UBXD1 as a potent p97 ATPase inhibitor and report structures of intact p97:UBXD1 complexes that reveal extensive UBXD1 contacts across p97 and an asymmetric remodeling of the hexamer. Conserved VIM, UBX, and PUB domains tether adjacent protomers while a connecting strand forms an N-terminal domain lariat with a helix wedged at the interprotomer interface. An additional VIM-connecting helix binds along the second AAA+ domain. Together these contacts split the hexamer into a ring-open conformation. Structures, mutagenesis, and comparisons to other adapters further reveal how adapters containing conserved p97-remodeling motifs regulate p97 ATPase activity and structure.

## INTRODUCTION

p97 (also called valosin-containing protein or VCP) is a AAA+ (ATPases Associated with diverse cellular Activities) molecular chaperone unfoldase with critical functions in many cellular processes including membrane fusion, chromatin remodeling, and protein homeostasis (proteostasis)^1,2^. Reflecting its critical roles in the cell, missense mutations in p97 cause multisystem proteinopathy (MSP, also called IBMPFD), amyotrophic lateral sclerosis, and vacuolar tauopathy, all characterized by defects in protein quality control and clearance pathways^3–5^. Additionally, p97 is under investigation as a cancer target due to its central roles in maintaining proteostasis, and is upregulated in many carcinomas^6,7^. The central mechanism of p97 that governs its diverse activities is the extraction of proteins from macromolecular complexes and membranes through hydrolysis-driven substrate translocation^8,9^. This process frequently facilitates substrate degradation by the 26S proteasome, as in the endoplasmic reticulum-associated degradation (ERAD) and ribosome quality control pathways^10,11^. In less well-understood pathways, p97 enables autophagic clearance of endosomes and damaged organelles such as lysosomes or mitochondria, potentially by regulatory remodeling of their surface proteomes^12–14^. p97 functions are directed by more than 30 adapter proteins that bind p97 to facilitate substrate delivery, control subcellular localization, and couple additional substrate processing functions such as deubiquitylation^15^. Thus, the many possible adapter interaction combinations may in part drive p97 functional diversity^16^. Some of these adapters are well-characterized, for example the ERAD-related UFD1/NPL4 and YOD1 recruit ubiquitylated substrates to p97 and catalyze ubiquitin chain removal, respectively^10^. However, the function of many adapters, such as the mitophagy- and lysophagy-related UBXD1, is unknown^12,17^.

p97 forms homohexamers that enclose a central channel through which unfolded proteins are threaded; each protomer consists of two AAA+ ATPase domains (D1 and D2), an N-terminal domain (NTD), and an unstructured C-terminal (CT) tail^18^. Initial structures of full-length or truncated p97 revealed a planar, symmetric hexamer in which the NTD adopts an ‘up’ conformation when D1 is bound to ATP, or a ‘down’ conformation co-planar with the ADP-bound D1 ring^18–24^. More recent substrate-bound structures revealed that the hexamer adopts a right-handed spiral with the conserved pore loops in D1 and D2 extending into the channel, contacting the substrate polypeptide in a manner similar to many AAA+ translocase complexes^8,9, 25–27^. Based on comparisons with related AAA+s, p97 may extract and unfold subtracts by a similar processive ‘hand over hand’ mechanism involving stepwise cycles of substrate binding and release^26–29^. However, given its diverse pathways and the central involvement of numerous adapters, alternate translocation paths may occur, potentially for processing substrates with complex topologies including those bearing branched ubiquitin chains^16^. Notably, substrate recognition and engagement are largely dependent on p97-adapter coordination^15^, yet with the exceptions of UFD1/NPL4^9,30,31^ and p37^32–34^, little is known about the mechanism by which adapters control p97 activity to enable these processes.

UBXD1 is a p97 adapter implicated in autophagic clearance of damaged mitochondria and lysosomes, among other functions^12,14,17,35,36^. MSP mutations impair UBXD1 binding and affect its associated cellular functions, demonstrating the importance of this adapter in multiple p97 pathways, but its function in p97 substrate processing is unknown^14,37,38^. UBXD1 contains three structurally defined p97 interaction motifs, but no known enzymatic or substrate interaction activities. A conserved VIM (VCP-interacting motif) helix interacts with the p97 NTD, while the PUB (peptide:*N*-glycanase and UBA or UBX-containing proteins) domain interacts with the CT tail^36,39–42^. Notably, UBXD1 contains a canonical UBX (ubiquitin regulatory X) domain that is also expected to bind the NTD. However, it is unclear how the VIM and UBX could function together given the overlapping binding sites. Moreover, the UBX domain is reported to not interact with p97 potentially due changes in binding pocket residues compared to other UBX domains^17,39,43^. Crystal structures of p97 truncations in complex with VIM, UBX, and PUB domains from other adapters have provided insight into isolated p97-adapter interactions, but how all these domains function together in an intact p97:UBXD1 complex is unknown. Furthermore, UBXD1 is the only known p97 adapter with both N- and C-terminal-interacting domains; this feature, coupled with the absence of substrate-binding domains and multiple NTD-interacting domains, raises key questions about the arrangement of UBXD1 on p97 hexamers, its binding stoichiometry, and effect on p97 structure and function.

To address these questions, we determined high-resolution cryo-EM structures of p97 in complex with full-length UBXD1. The structures reveal multiple stoichiometric arrangements of UBXD1 that involve extensive contacts with p97 via its VIM, PUB, and UBX domains, as well as previously uncharacterized helices. Remarkably, in singly-bound states, UBXD1 interacts across three p97 protomers, dramatically remodeling the planar p97 hexamer into a partial spiral, open-ring state in which the separated protomers remain tethered by UBXD1. A lariat structure wraps around an NTD with two helices interacting at the protomer interface, functioning as a wedge that displaces D1 interprotomer contacts, while a short helix adjacent to the VIM alters the D2 conformation. These UBXD1 interactions coincide with a potent loss of p97 ATPase activity. Together with mutagenesis and additional structural characterization this work reveals distinct roles for the UBXD1 domains and identifies how a network of protein-protein adapter interactions coordinate to remodel p97 structure and control function.

## RESULTS

### Structures of p97:UBXD1 reveal extensive interactions and hexamer remodeling

In addition to the canonical VIM, UBX and PUB domains, UBXD1 is also identified to contain transiently structured helices, here referred to as H1/H2 (residues 1-25) and H4 (residues 75-93), that weakly interact with p97^36,37^ as well as additional uncharacterized regions (residues 278-334). To further investigate the predicted structure of full-length UBXD1, we analyzed the AlphaFold model^44,45^, which revealed structures for the canonical domains, H1/H2, H4 and an additional four-helical element between the PUB and UBX, while the remaining sequence appears unstructured (Fig. 1a,b). The four-helical region hereinafter is referred to as the helical lariat based on our structural analysis and homology to the ASPL adapter (see below)^46^. Notably, this AlphaFold analysis indeed predicts structures for uncharacterized regions across UBXD1 that, in addition to the VIM, UBX, and PUB, may interact with p97 and contribute to UBXD1 function.

**Fig 1.**
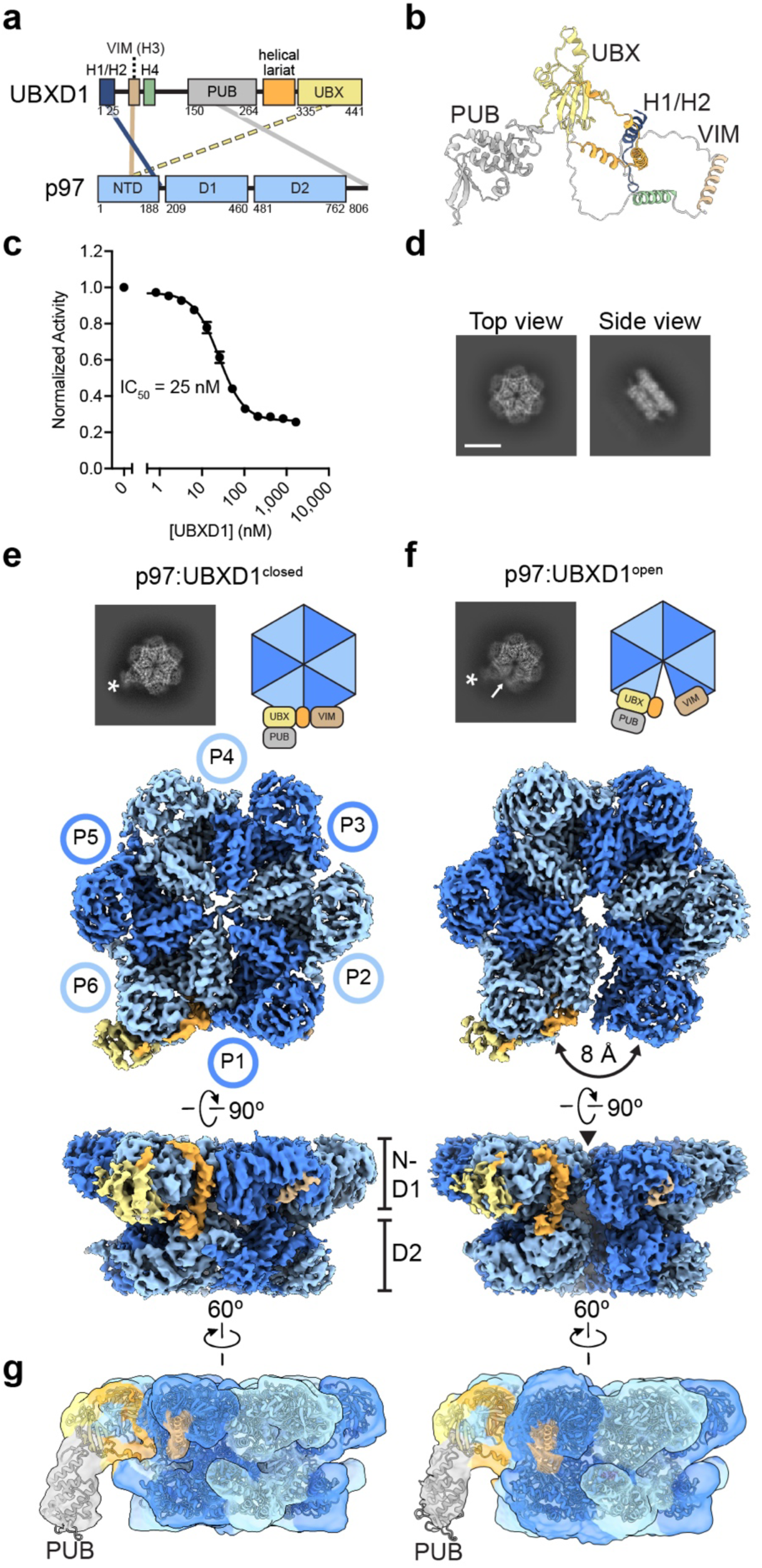
Cryo-EM structures of p97:UBXD1 closed and open states. (**a**) Domain schematics of UBXD1 and p97 (not to scale) showing reported interactions (solid lines) between conserved domains^36,39–41^ and the canonical UBX-NTD interaction previously reported to not occur for UBXD1 (dashed line)^17,39,43^. (**b**) AlphaFold model of UBXD1 showing structured regions (H1/H2, VIM, H4, PUB, helical lariat and UBX) colored as in (**a**). (**c**) Steady state ATPase activity (Y-axis, normalized to activity at 0 nM UBXD1) of p97 at increasing concentrations of UBXD1 (X-axis), resulting in a calculated IC_50_ of 25 nM. Error bars represent standard deviation and data are from three independently performed experiments. (**d**) Representative 2D class averages following initial classification of the full p97:UBXD1 dataset, showing p97 hexamer and no additional density for UBXD1 (scale bar equals 100 Å). Final cryo-EM reconstructions of (**e**) p97:UBXD1^closed^ and (**f**) p97:UBXD1^open^ states with top-view 2D projections showing UBX/PUB density (*) and open p97 ring (arrow) compared to cartoon depiction of p97:UBXD1 complex (top row); (below) cryo-EM density maps (p97:UBXD1^open^ is a composite map, see methods), colored to show the p97 hexamer (light and dark blue, with protomers labeled P1-P6) and UBXD1 density for the VIM (brown), UBX (yellow) and lariat (orange) domains. The 8 Å separation between protomers P1 and P6 is indicated for p97:UBXD1^open^. (**g**) Low-pass filtered maps and fitted models of p97:UBXD1^closed^ (left) and p97:UBXD1^open^ (right) exhibiting low resolution density for the PUB domain (gray).

To analyze the p97-UBXD1 interaction, we first determined the effect of UBXD1 on p97 ATPase activity using full-length, wildtype, human proteins (Fig. 1c). This revealed a potent inhibitory activity, with an IC_50_ of ~25 nM for UBXD1. We next analyzed the nucleotide dependence of the p97-UBXD1 interaction, as many adapters exhibit nucleotide binding specificity to p97 that is related to the up/down conformational plasticity of the NTDs^37,47,48^. By analytical size exclusion chromatography (SEC) we identify that following incubation with either ADP or the slowly hydrolysable ATP analog ATPγS, UBXD1 co-elutes with p97 in fractions slightly shifted from the hexamer peak, indicating complex formation under both nucleotide conditions (Extended Data Fig. 1a,b). A modest shift in elution is identified with ADP compared to ATPγS, however both nucleotide-bound states appear to support UBXD1 binding under these conditions, indicating UBXD1 may interact irrespective of the NTD state. Nonetheless, based on the reduced band intensity of UBXD1 compared to p97, UBXD1 binding appears to be sub-stoichiometric with respect to p97 hexamer despite incubations at high concentration (20 and 10 µM, respectively). A previous study reported UBXD1 binds p97 sub-stoichiometrically, but with a modest affinity (*K*_D_ ~3.5 µM), consistent with our SEC results^41^. However, we predict a much higher binding affinity based on the ~25 nM IC_50_ for ATPase inhibition we identify (Fig. 1c).

Based on these results and the reported increased NTD stability in the ADP state^19^, p97:UBXD1 cryo-EM samples were first prepared with saturating concentrations of ADP (5 mM). Reference-free two-dimensional (2D) class averages show well-defined top and side views of the p97 hexamer with the NTDs in the down conformation and co-planar with the D1 ring, as expected based on the presence of ADP (Fig. 1d and Extended Data Fig. 1c). However, additional density around the hexamer was not identified in these initial classifications. Considering potential UBXD1 flexibility and binding heterogeneity could impact 2D analysis, rounds of three-dimensional (3D) classification were performed next with a large (~5.5 million particle) dataset in order to better classify potential bound states. Following an initial classification, three well-defined states were identified. Classes 1 and 2 contained additional globular density likely corresponding to the UBX domain of UBXD1 while class 3 appeared as a well-resolved p97 hexamer without the additional globular density (Extended Data Fig. 1d). Subsequent classification and refinement of the UBX-containing states enabled high-resolution structure determination (3.3-3.5 Å) of p97:UBXD1 in two configurations termed p97:UBXD1^closed^ and p97:UBXD1^open^ (Fig. 1e,f and Table 1). Individual p97 protomers are denoted counterclockwise P1-P6 based on asymmetry of p97:UBXD1^open^. Additional UBXD1 binding configurations were identified in the classification and further characterized (see below and Extended Data Fig. 1d). Notably, the singly-bound closed and open states exhibit the most p97 conformational changes and extensive contacts by UBXD1 and thus are largely the focus of this study. Analysis of the closed and open states revealed that the protomers unbound to UBXD1 were well-resolved, while protomers P1 and P6 contain additional density corresponding to UBXD1 and comprise a flexible seam that in the open state is more poorly resolved relative to the rest of the hexamer (Fig. 1e,f). Therefore, focused classification and local refinement of P1 and P6 in the open state was performed separately to improve the resolution of these regions, and a composite map was generated (Extended Data Fig. 1d and methods). High-resolution map features in both the closed and open states enabled building of atomic models for the p97 hexamer, and the UBXD1 domains predicted to be structured by the AlphaFold model are all identified except H1/H2 and H4 (Fig. 1e,f, Extended Data Fig. 2a-d,k, and Supplementary Video 1).

**Table 1.**
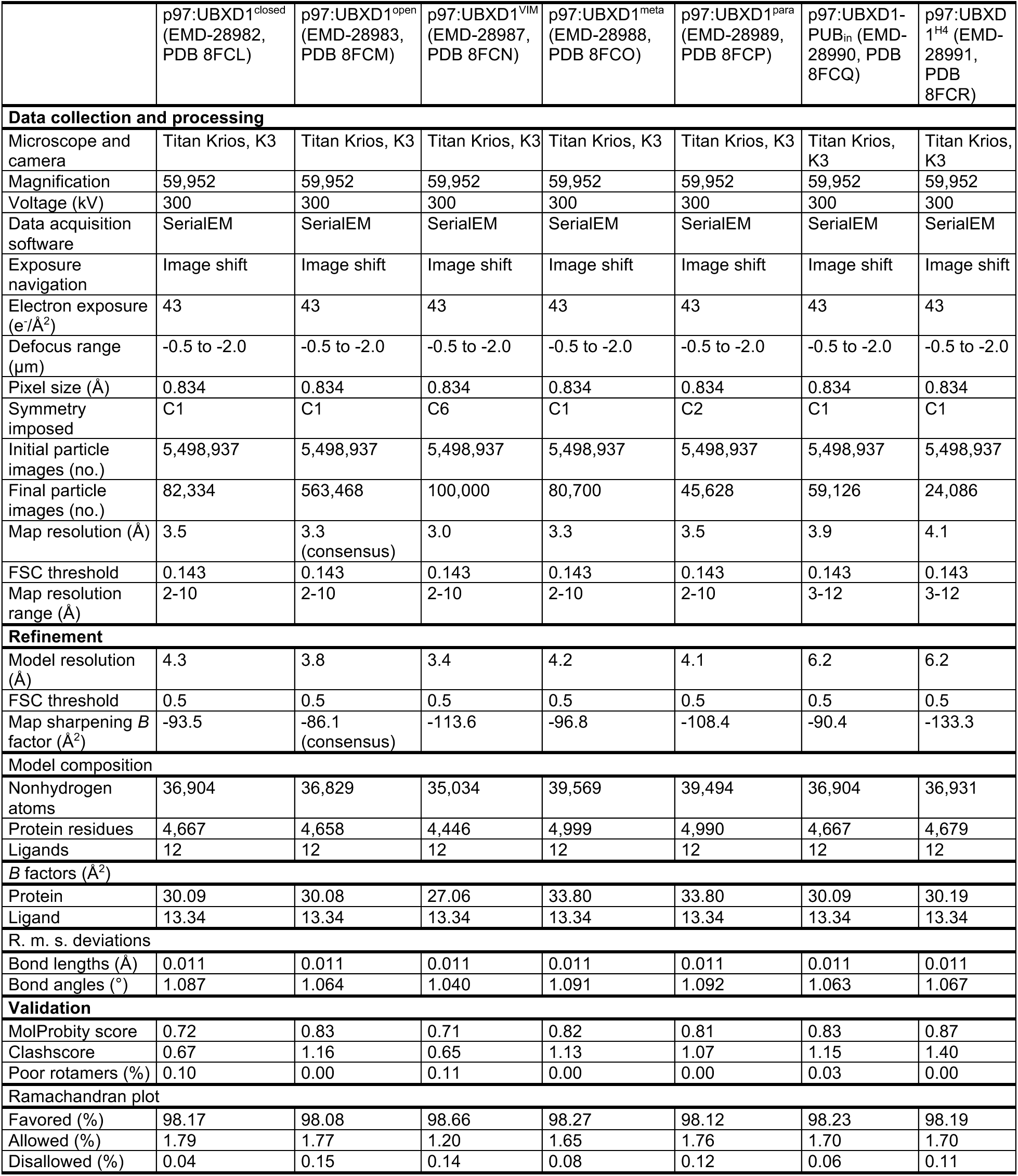
Cryo-EM data collection, refinement, and validation statistics of structures from the p97/UBXD1^WT^/ADP dataset.

The p97 domains are well resolved in both structures with the NTDs adopting the down conformation and density corresponding to bound ADP is present in all D1 and D2 nucleotide pockets and coordinated by conserved AAA+ residues (Extended Data Fig. 1e-g). In the p97:UBXD1^closed^ structure the D1 and D2 are in a planar, double ring conformation and one molecule of UBXD1 is identified, binding across protomers P1 and P6. For the p97:UBXD1^open^ structure a single UBXD1 is similarly bound but the protomer-protomer interface between P1 and P6 is separated by ~8 Å relative to p97:UBXD1^closed^, creating an open-ring arrangement of the p97 hexamer. For both structures, strong density corresponding to the VIM and UBX domains is observed, as well as for two previously uncharacterized helices and an associated N-terminal linker (Fig. 1e,f). Together, these helices and linker encircle the NTD of the P6 protomer, forming a helical lariat (discussed below). At lower density threshold, the PUB domain becomes apparent and is positioned below the UBX, adjacent to the p97 D2 ring, but appears to make no substantial contact with the hexamer, likely resulting in significant flexibility and explaining the lower resolution (Fig. 1g). Weak density resembling the VIM is also present in the NTDs of P2-P5 in both the closed and open states, indicating partial binding to these protomers, which is likely a consequence of protein concentrations needed for cryo-EM (Extended Data Fig. 1h). Overall, these structures indicate that UBXD1 binding drives conformational changes and ring opening at a protomer interface that is tethered by multi-domain interactions involving the VIM, UBX, and a lariat structure across the two seam protomers.

### Additional structures identify alternate UBXD1 binding stoichiometries and nucleotide-dependent hexamer remodeling

During 3D classification additional structures with different UBXD1 configurations were identified (Extended Data Fig. 1d,i, 2, and Table 1). A prevalent class in the initial classification, termed p97:UBXD1^VIM^, closely resembles the structure of the planar ADP-bound p97 (Cα root-mean-square deviation (RMSD) = 1.0 Å)^19^, but features additional helical density in the NTD cleft of all protomers that corresponds to the UBXD1 VIM (Extended Data Fig. 1i). Similar to P2 through P5 in the closed and open states, we surmise that under our *in vitro* conditions VIM binding predominates for this class, potentially displacing the UBX and other contacts. Notably, given the identical hexamer arrangement compared to p97^ADP^, we conclude that VIM interactions on their own are insufficient to induce conformational rearrangements in p97. Subclassification additionally revealed two more minor states, termed p97:UBXD1^meta^ and p97:UBXD1^para^, in which density for the VIM, PUB, lariat, and UBX are also observed at other positions (across P2-P3 or P3-P4, respectively) in the p97 hexamer, in conformations similar to p97:UBXD1^closed^ (Extended Data Fig. 1d,i). These structures reveal that two UBXD1 molecules can bind the hexamer with productive VIM, UBX, and lariat interactions when properly spaced to allow for binding across two adjacent protomers. Notably, for all classifications the p97 open ring is only observed in the singly-bound UBXD1 configuration, indicating that this conformation is specifically driven by one fully-bound UBXD1 per p97 hexamer.

We identify UBXD1 also binds p97 in the presence of ATPγS, but with a reduced shift in elution volume compared to ADP (Extended Data Fig. 1a,b). Therefore, we next sought to determine the extent of UBXD1-mediated structural remodeling of p97 in its ATP state. Cryo-EM analysis revealed three predominant classes: a UBXD1-bound hexamer closely resembling the closed state from the ADP dataset, a symmetric hexamer with NTDs in the up, ATP conformation and no density for UBXD1, and a symmetric hexamer with ‘down’ NTDs and VIM density, similar to the VIM-only state from the ADP dataset (Extended Data Fig. 3a). Notably, no state analogous to the open p97:UBXD1 complex was identified. Refinement of these states allowed for the unambiguous assignment of ATPγS for density in all nucleotide-binding pockets (Extended Data Fig. 3b,c). The observation of ‘down’ NTDs in ATPγS-bound D1 domains is striking, indicating that UBXD1 interactions may override nucleotide-promoted NTD conformation, a finding not previously described in intact complexes. Indeed, the NTD-down arrangement is identified even in the VIM-only state with ATPγS, suggesting that interaction by the UBXD1 VIM alone is sufficient to regulate p97 NTD conformation. In sum, these results suggest that while UBXD1 readily binds p97 in the ATP state, the interactions are insufficient to promote full ring opening, as observed with ADP. The absence of an open state may be due to increased inter-protomer interactions and hexamer stability, such as through trans-arginine finger contacts with the γ-phosphate. Indeed, in substrate-bound AAA+ complexes, hydrolysis at the spiral seem is thought to destabilize the interprotomer interface, facilitating substrate release and rebinding during stepwise translocation^49^. Our findings therefore indicate that UBXD1 may function during the p97 catalytic cycle by promoting ring opening specifically in a post-hydrolysis state.

### UBXD1 drives large D1-D2 conformational changes that open the p97 hexamer ring

To identify conformational changes in ADP-bound p97 complexes that are driven by UBXD1 binding, the p97:UBXD1^closed^ and p97:UBXD1^open^ structures were aligned to the previously published structure of ADP-bound p97^19^. RMSD values reveal extensive changes across the D1 and D2 for the seam protomers (P1 and P6) in both UBXD1-bound states (Fig. 2a,b and Extended Data Fig. 4a,b). While P1 and P6 are similarly rotated away from each other in both states, the magnitudes of these displacements are larger in the open state, resulting in the observed disruption of the protomer interface. This indicates that these states are intermediates in the same conformational path of hexamer opening (Supplementary Video 2). Additionally, while the hexamer in the closed state is planar and resembles substrate-free structures, the open state has a right-handed spiral with an overall elevation change of 7 Å (Fig. 2c). This is largely due to a significant 9° downward rotation of P1, which has the largest RMSD values of any protomer in either state (Fig. 2c). In addition to the movements of entire protomers, there is a notable rotation between the small and large subdomains of the D2 in protomer P1 in both the closed and open states (Fig. 2d). This rotation is particularly evident in the open state, where the small subdomain is rotated upward by 10° relative to p97^ADP^ and 12° relative to p97:UBXD1^closed^ (Fig. 2d). This rotation, and the separation of P1 and P6, causes a small helix (α5’)^18,24^ mediating contact between the D2 domains of the seam protomers to disappear from the density map, likely as a result of increased flexibility; this helix is normally positioned on top of α12’ of the D2 domain of the counterclockwise protomer (Extended Data Fig. 4c and Fig. 2b).

**Fig. 2.**
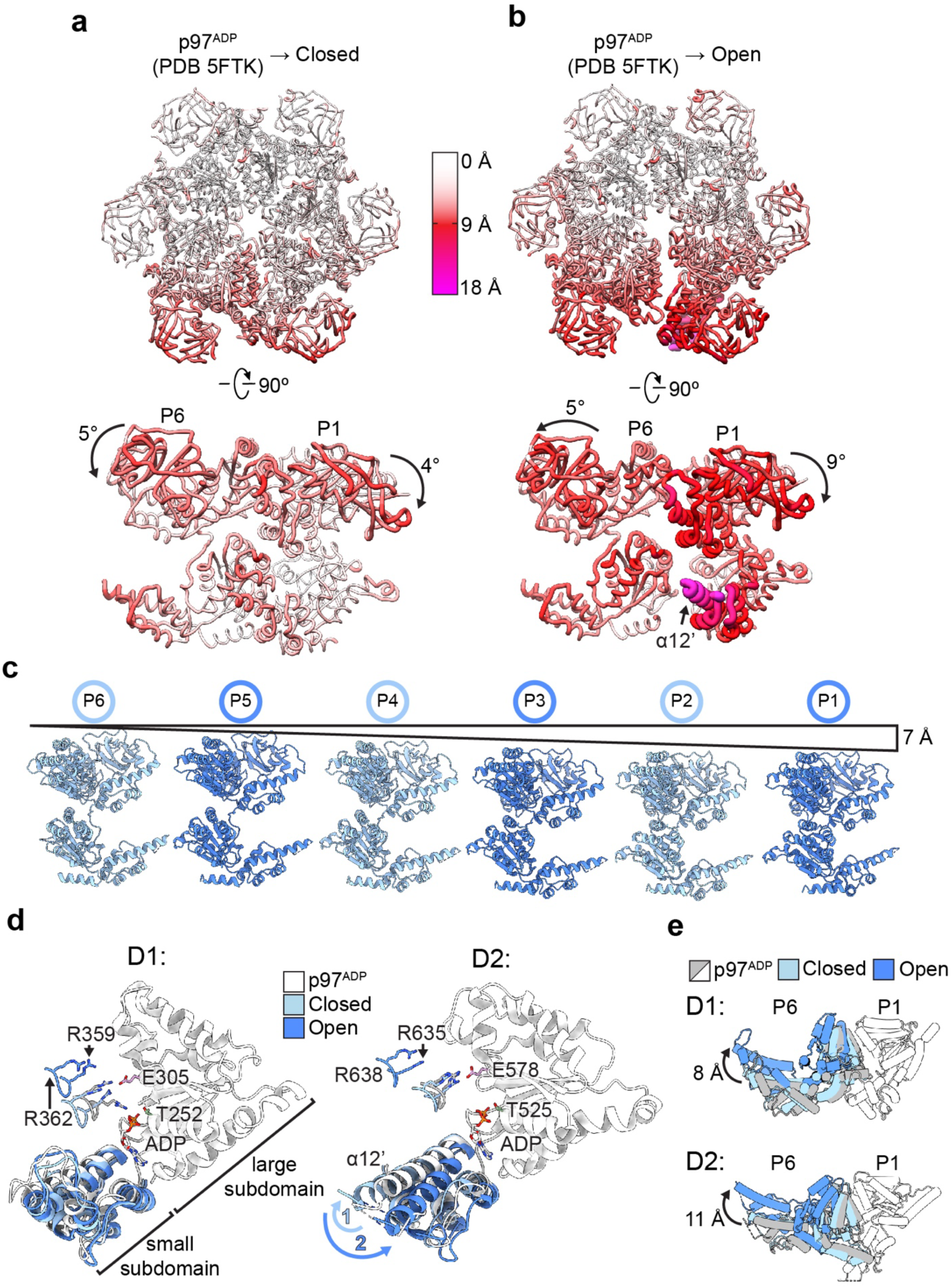
UBXD1-mediated p97 hexamer remodeling. The p97 hexamer and rotated side view of the seam protomers P1 and P6 for (**a**) p97:UBXD1^closed^ and (**b**) p97:UBXD1^open^ structures, colored according to Cα RMSD values relative to the p97^ADP^ symmetric state (PDB 5FTK, aligned to P3 and P4). The largest changes (>15 Å, magenta with wider tubes) are identified for P1 α12’ (arrow) in the open state, intermediate changes (~10 Å, red) for P1 and P6 with rotations of the NTDs relative to p97^ADP^ shown, and small/no changes for the remaining regions (<5 Å, white). (**c**) Side by side view of individual protomers aligned based on position in the p97:UBXD1^open^ hexamer, showing vertical displacement along the pseudo-C6 symmetry axis. (**d**) Overlay of the D1 (left) and D2 (right) AAA+ domains of P1 for p97^ADP^, p97:UBXD1^closed^, and p97:UBXD1^open^, aligned to the large subdomains and colored as indicated. ADP is shown with conserved Walker A/B and trans-acting (P6) Arg finger residues indicated. The large rotation of the D2 small subdomain, exemplified by α12’, is shown (relative to p97^ADP^) for the closed (1) and open (2) states. (**e**) Top view overlay of the D1 (upper) and D2 (lower) domains for the P6-P1 pair in the three states and aligned to P1 to show relative rotations of P6, colored as indicated. Rotations shown are from the p97^ADP^ to the p97:UBXD1^open^ state and determined from centroid positions of the D1 and D2 domains.

Notably, the P1-P6 conformational changes are different across the D1 and D2 domains (Fig. 2e). In the closed state the D1 domains of P1 and P6 are separated by 3 Å relative to p97^ADP^, while there is negligible D2 separation. In the open state the D1 and D2 domains are separated by 8 Å and 11 Å, respectively. These changes dramatically remodel the nucleotide binding pockets of P1 by displacing the arginine fingers from P6 (Fig. 2d). These separations likely preclude ATP hydrolysis in P1, which could explain the ATPase inhibition observed biochemically (Fig. 1c). Additionally, due to the expansion of the P1-P6 interface in the open state, contacts between other protomers are compressed: the average rotation between adjacent protomers is 57°, or 3° smaller than the angle in a perfectly symmetric hexamer. These compressions could potentially cause subtle deformations of nucleotide binding pocket geometry, which could also impair hydrolysis. Finally, to confirm the spiral architecture of the open state, 3D variability analysis was performed jointly on particles from the closed and open states (Supplementary Video 3). This reveals the transition from a planar UBXD1-bound hexamer to a spiral, indicating that the closed and open states are likely in equilibrium and UBXD1 binding splits the p97 hexamer at P1-P6, causing all protomers to rotate along the hexamer C6 symmetry axis.

### Canonical interactions by the VIM, UBX and PUB indicate binding across three protomers of p97

The p97:UBXD1^closed^ and p97:UBXD1^open^ structures reveal high-resolution views of p97 bound to an adapter containing conserved VIM, UBX and PUB domains, revealing how these domains together coordinate interactions across the hexamer. In contrast to previous binding studies^17,39,43^, both the VIM and UBX interact with p97, making canonical interactions with adjacent NTDs (Fig. 3). The 18-residue VIM helix is positioned in the NTD cleft of P1, similar to structures of isolated domains, but is distal to other UBXD1 density, and comprises the only major contact with the P1 protomer (Fig. 3a,b and Extended Data Fig. 5a,b). Notably, a conserved Arg residue (R62) required for p97 binding projects into the NTD in a manner similar to that of other NTD-VIM complexes, potentially forming a salt bridge with D35 of the NTD and a hydrogen bond with the backbone carbonyl of A142^41^. The VIM appears anchored at its N-terminus by an additional salt bridge between E51 of UBXD1 and K109 of the NTD and by hydrogen bonding between the backbone carbonyl of E51 and Y143 of the NTD. Additional hydrophobic contacts along the VIM could further stabilize this interaction (Fig. 3b, right panel).

**Fig. 3.**
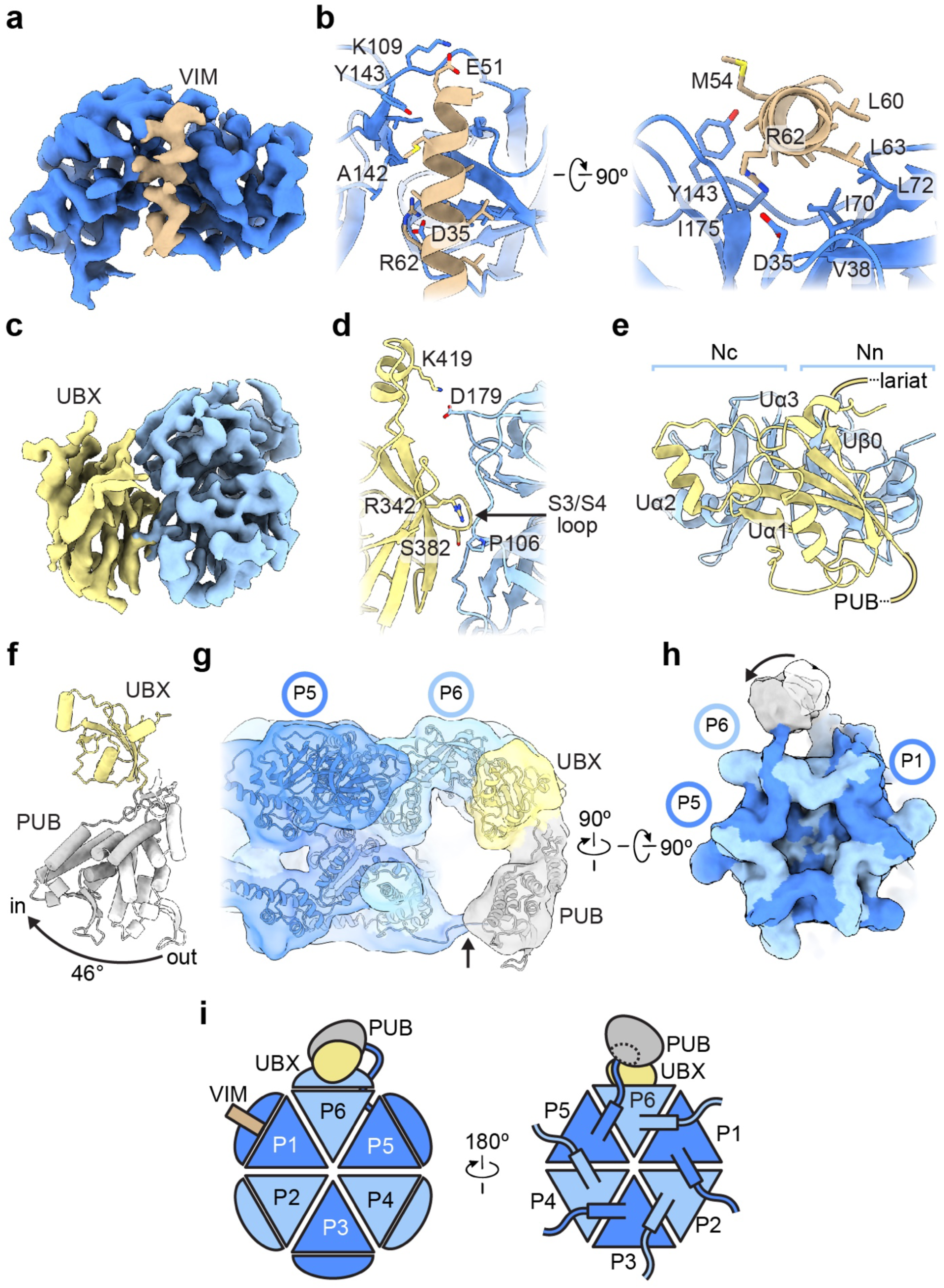
Interactions by conserved VIM, UBX and PUB domains of UBXD1 across the p97 hexamer. (**a**) Sharpened map of P1 NTD (dark blue) and the VIM helix (brown) from the p97:UBXD1^closed^ structure. (**b**) Model showing VIM helix interactions with the NTD, colored as in (**a**) with labeled interacting residues. (**c**) Sharpened map of the P6 NTD (light blue) and UBX domain (yellow). The model of the UBX and NTD in p97:UBXD1^closed^ is shown (**d**) depicting UBX-NTD contacts, including the conserved S3/S4 loop contact^59^ (arrow) and (**e**) with non-canonical structural elements Uα2, Uα3, and Uβ0, colored as in (**c**). (**f**) Overlay of PUB from p97:UBXD1-PUB_in_ (gray) and p97:UBXD1-PUB_out_ (white) (see methods), aligned to the UBX (yellow) domain, showing 46° rotation of the PUB domain position. (**g**) Low-pass filtered map and model of p97:UBXD1-PUB_in_ depicting PUB domain contact with p97 and model for C-terminal HbYX tail interaction from the adjacent P5 protomer (arrow) and (h) bottom view of the hexamer map with out (white) and in (gray) positions of the PUB. (**i**) Cartoon of p97:UBXD1^closed^ depicting UBXD1 interactions across three p97 protomers (P1:VIM, P6:UBX, and P5:PUB) through canonical p97-interacting domains.

The UBX domain is bound to the NTD of the clockwise protomer (P6) in a manner similar to UBX-NTD structures from other adapter proteins (Fig. 3c,d and Extended Data Fig. 5c,d). This is surprising because the canonical Phe-Pro-Arg p97-interacting motif located on the S3/S4 loop is replaced by Ser-Gly-Gly in UBXD1, thus eliminating electrostatic and hydrophobic interactions identified in other structures^15^. However, another Arg residue important for interaction with p97, R342, is conserved in UBXD1, and likely hydrogen bonds with the backbone carbonyl of P106. While the UBX in this structure contains the canonical β-grasp fold characteristic of all UBX domains, it features an additional β-strand (Uβ0) proximal to the N-terminal lobe of the NTD (Nn) that connects the PUB to the UBX to the helical lariat (Fig. 3e). Additionally, a C-terminal extension consisting of two alpha helices (UBX helices 2 and 3, hereafter referred to as Uα2 and Uα3) connected by unstructured linkers is positioned on the apical surface of the canonical UBX and wraps over the β-sheet. Finally, a potential salt bridge between K419 of Uα2 and D179 of the NTD could also stabilize the UBX-NTD interaction.

The PUB domain binds the HbYX (hydrophobic, Tyr, any amino acid) motif located at the end of the flexible p97 CT tail^40^. Density for this domain is more poorly resolved in both the closed and open structures, which prompted us to perform focused classification of this region (Extended Data Fig. 1d). Two resulting classes show improved definition for the PUB, enabling the AlphaFold model for this region to be fit unambiguously into the density (Extended Data Fig. 5e,f). In class 1 (hereafter referred to as p97:UBXD1-PUB_out_), the PUB domain is positioned similarly to that in the closed model, projecting straight downward from the UBX domain. In class 2 (hereafter referred to as p97:UBXD1-PUB_in_) the PUB domain is rotated 46° about a linker connecting the PUB and UBX domains, and points towards P6 (Fig. 3f, Extended Data Fig. 2, 5f, and Table 1). In this class strong connecting density is observed between the PUB and the bottom surface of P6, indicative of binding to a p97 CT tail (Fig. 3g,h). Notably, these tails project across the base of the adjacent counterclockwise protomer, such that UBXD1 density on P6 is closest to the P5 tail, not that of P6. Thus, the position of the PUB domain in p97:UBXD1-PUB_in_ indicates binding to the P5 tail (Fig. 3g,h). Inspection of the p97:UBXD1-PUB_out_ map reveals weak density suggestive of a similar P5-PUB interaction, indicating that the CT tail may remain bound in multiple PUB conformations (Extended Data Fig. 5h-k), though the strongest density is present in p97:UBXD1-PUB_in_. In sum, a single molecule of UBXD1 appears to interact across three p97 protomers (P1, P5, and P6) simultaneously through interactions by the VIM, UBX and PUB domains (Fig. 3i). These extensive interactions with both faces of the p97 hexamer are a remarkable feature of UBXD1, and are unique among all adapters currently structurally characterized.

### The UBXD1 helical lariat and H4 make distinct p97 D1-D2 interprotomer interactions

The UBXD1 helical lariat is among the most striking structural features of the p97:UBXD1 complexes due to its intimate interaction with all three domains of the P6 protomer (Fig. 4a-e and Extended Data Fig. 6a). Based on the AlphaFold prediction and what is resolved in our structures, it is composed of four helices (hereafter referred to as Lα1-4) that are inserted between Uβ0 and Uβ1 of the UBX domain. Together, these helices completely encircle the P6 NTD (Fig. 4a). Lα1 and Lα2 are positioned along the top of the P6 NTD, while the longer Lα3 and Lα4 helices bind at the P6-P1 protomer interface. Lα3 is situated at the D1 interface, and makes numerous electrostatic contacts with residues in both the N-terminal and D1 domains of P6, as well as minor hydrophobic contacts with the D1 domain of P1 (Fig. 4b,c). Lα4 is positioned proximal to the D2 domain, and connects back to the UBX domain, completing the lariat. A salt bridge involving K325 of Lα4 and E498 of the D2 domain is the only significant contact this helix makes with p97. Lα2 also contacts the NTD using two Phe residues (F292 and F293) that project into the NTD, making van der Waals contacts with K62, V99, and R25 (Fig. 4d). A short loop connects Lα3 and Lα4, and anchors the lariat into the D2 domain using residues L317 and T319 (Fig. 4b,e). Additionally, the Lα3-Lα4 arrangement is stabilized by a tripartite electrostatic network involving R313 of Lα3, R318 of the Lα3-Lα4 loop, and E326 of Lα4 (Fig. 4e). Considering the interaction of Lα3 along the D1 domain of P6 displaces typical D1-D1 contacts between P1 and P6, we postulate these contacts likely contribute substantially to the D1 conformational changes and ring-opening identified in the closed and open states of p97. Interestingly, the binding site of the linker between Lα3 and Lα4 overlaps with that of the p97 allosteric inhibitor UPCDC30245^19^, suggesting that occupancy of this site is a productive means to alter p97 function (Extended Data Fig. 6b).

**Fig. 4.**
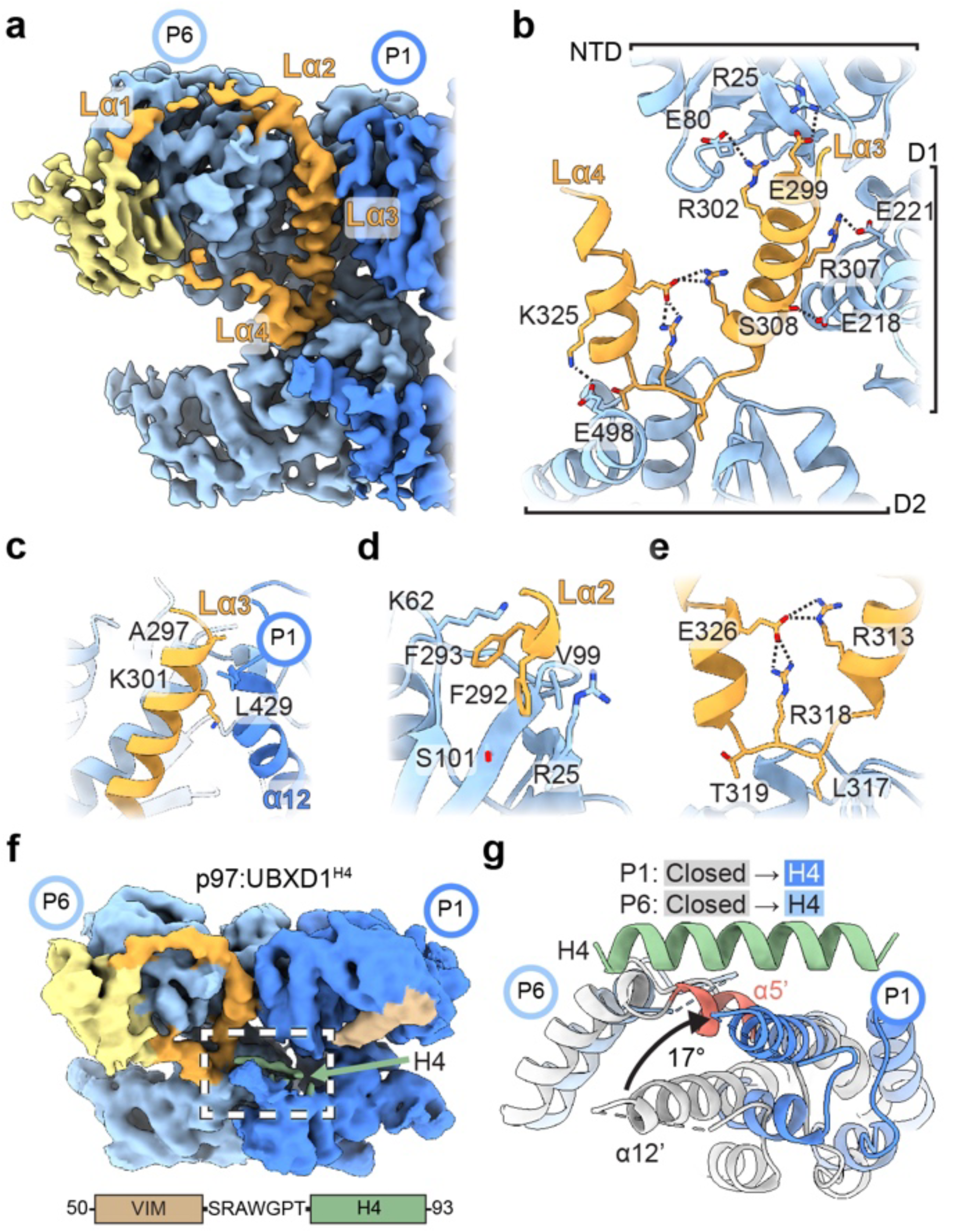
p97 remodeling interactions by UBXD1 helical lariat and VIM-H4. (**a**) Closed state map (from p97:UBXD1^meta^) showing density for the UBXD1 helical lariat (orange) and UBX (yellow) encircling the P6 NTD with Lα2, Lα3, and Lα4 interacting along the P6-P1 interprotomer interface. (**b**) Expanded view showing Lα3 and Lα4 (orange) contacts with P6 across the NTD, D1, and D2, including putative electrostatic interactions (dashed lines). (**c**) View of the P6-P1 interface showing key contacts by Lα3 with the D1 α12 helix of protomer P1. View of (d) Lα2 interactions involving hydrophobic packing into the NTD and (**e**) Lα3 and Lα4 intra-lariat contacts and contacts with D2, stabilizing the helical lariat. (**f**) Unsharpened map of p97:UBXD1^H4^, showing density for H4 (green) adjacent to the VIM (brown) and along the P6-P1 interface. Shown below is an expanded view of the VIM-H4 sequence, featuring only a short 7-amino acid linker connecting the two helices. (**g**) Modeled view (see Extended Data Fig. 6c) of helix H4 interacting across the D2 domains at the P1-P6 interface with p97 from p97:UBXD1^H4^ (P1: dark blue, P6: light blue), and p97:UBXD1^closed^ (gray) overlaid (by alignment of the P1 D2 large subdomain) to show conformational changes at the P6-P1 interface including displacement of P6 helix α5’ (red) and large rotation of P1 α12’ in p97:UBXD1^H4^.

Further classification of p97:UBXD1^closed^, which was chosen due to the reduced flexibility compared to the open state, was performed to potentially resolve additional regions reported to interact with p97^36,37^, including H1/H2 and H4 (Extended Data Fig. 1d). This analysis revealed an additional state with an overall similar conformation to p97:UBXD1^closed^, but features low resolution density on top of the D2 domain of the P1 protomer (Fig. 4f, Extended Data Fig. 2, and Table 1). This region likely corresponds to H4 because of its proximity to the C-terminus of the VIM, which is predicted to be connected to H4 by a 7-residue linker (Fig. 1b). Indeed, the H4 helix docks well into this density, albeit the resolution was not sufficient for its precise orientation (Extended Data Fig. 6c). In this structure (p97:UBXD1^H4^), H4 binding is associated with an upward rigid body rotation of the D2 small subdomain by ~17° relative to the analogous domain in the closed state (Fig. 4g and Supplementary Video 4). This upward rotation displaces a short helix (α5’) from the D2 domain of P6, and therefore breaks D2-D2 contacts between the seam protomers. As in p97:UBXD1^open^, α5’ is not present in the density map, likely due to increased flexibility (Extended Data Fig. 6d). Notably, a similar rotation of the D2 small subdomain of P1 is identified in p97:UBXD1^open^, supporting potential H4 occupancy (Fig. 2e). This prompted further inspection of the experimental density for this state. Indeed, weak density positioned atop the D2 domain was identified by examining the open state map at a low threshold, in a similar position as in p97:UBXD1^H4^ (Extended Data Fig. 6e). This indicates that H4 may be associated with the hexamer during ring opening, and places p97:UBXD1^H4^ as an intermediate between the closed and open states. Based on this analysis, we predict that H4 interactions play a key role in weakening the D2 interprotomer contacts, thereby driving localized opening of the D2 ring.

### The helical lariat and H4 are conserved p97-remodeling motifs

Given the striking rearrangements of the p97 hexamer driven by UBXD1, efforts were undertaken to identify other p97 adapters with helical lariat or VIM-H4 motifs that could similarly remodel p97 contacts. To this end, Dali searches^50^ against all structures in the Protein Data Bank and against the AlphaFold database were performed, first using the UBXD1 UBX-helical lariat structure as an input. This search revealed one protein, alveolar soft part sarcoma locus (ASPL, also called TUG or UBXD9), with a highly similar UBX-helical lariat arrangement (Fig. 5a and Extended Data Fig. 7a). Comparison of p97:UBXD1 structures determined here to structures of a heterotetramer of a truncated ASPL construct bound to p97^46^ reveal a conserved interaction with p97 (Fig. 5b). The FF motif in Lα2, as well as the highly charged nature of the helices corresponding to Lα3 and Lα4, are conserved in ASPL (Extended Data Fig. 7a). Intriguingly, ASPL also inhibits p97 ATPase activity, and has been demonstrated to completely disassemble p97 hexamers into smaller oligomers and monomers^46,51^. However, the construct used for structure determination lacks several other p97-interacting domains, leaving unclear the effect on hexamer remodeling in the context of the full-length protein. While we find no evidence of a similar hexamer disruption in UBXD1 based on our SEC or cryo-EM analysis (Extended Data Fig. 1a-c), the split ring of the p97:UBXD1^open^ structure is compelling as a related function of the UBX-helical lariat in the context of UBXD1 with its additional p97 binding domains.

**Fig. 5.**
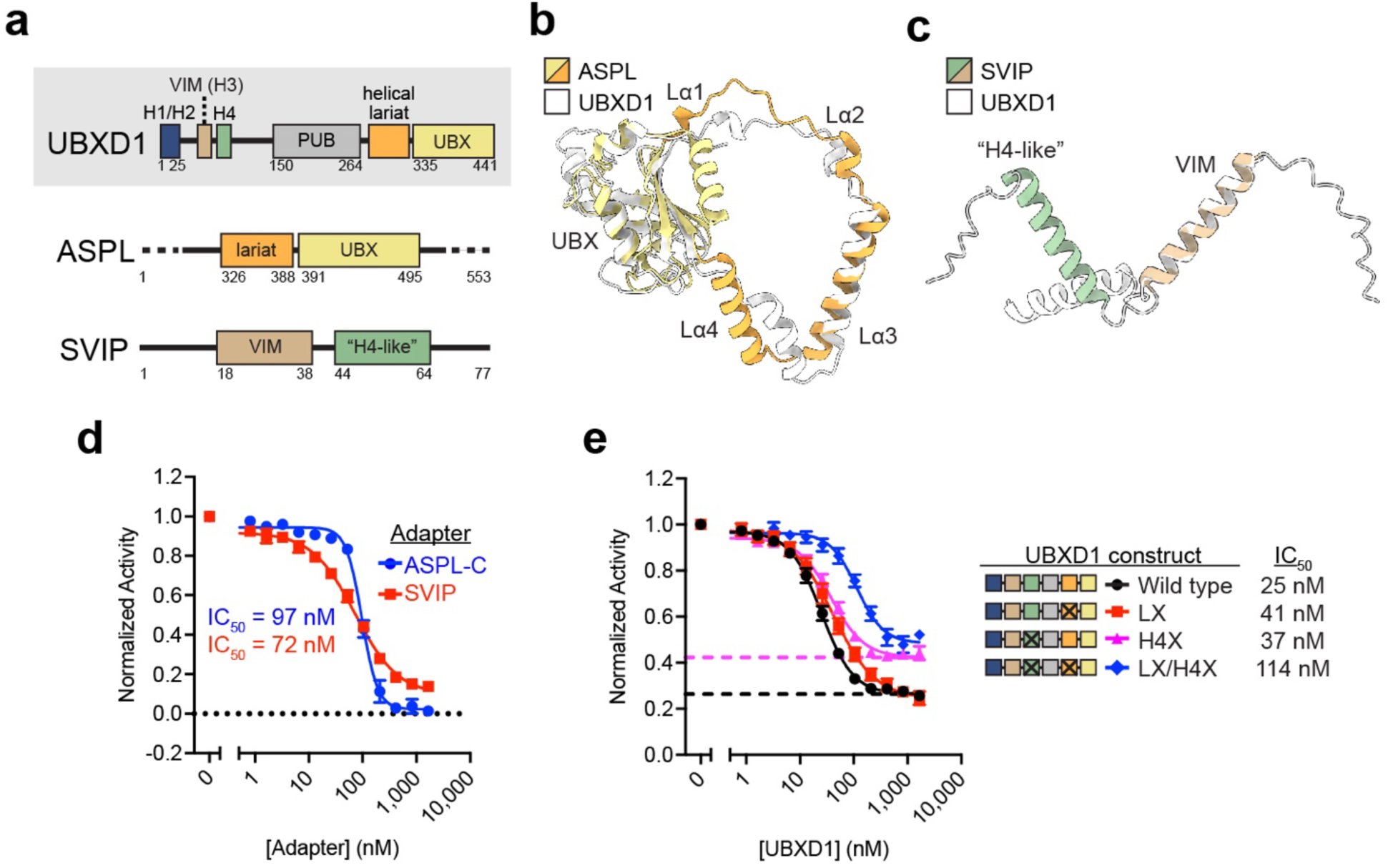
Analysis of the helical lariat and VIM-H4 as conserved p97-remodeling motifs. (**a**) Domain schematics of UBXD1, ASPL, and SVIP (not to scale). Overlay of the (**b**) UBX-helical lariat of ASPL (residues 318-495 from PDB 5IFS, colored as in (**a**)) and UBXD1 (residues 270-441 from the p97:UBXD1^closed^ model, in white) and the (**c**) VIM-“H4-like” region of SVIP (AlphaFold model, colored as in (**a**)) and UBXD1 (residues 50-93 from the AlphaFold model, in white). (**d**) Steady state ATPase activity of p97 as a function of ASPL-C or SVIP concentration. Error bars represent standard deviation and data are from three independently performed experiments. Calculated IC_50_s are shown below. (**e**) Steady state ATPase activity of p97 as a function of UBXD1 protein concentration for WT, LX (lariat mutant: E299R/R302E/R307E/E312R), or H4X (helix H4 sequence scramble); maximum concentration tested was 1.67 µM. Dashed lines represent the minimal activity (or maximal UBXD1 inhibition) obtained from the corresponding curve fit. Error bars represent standard deviation and data are from three independently performed experiments. Calculated IC_50_s and schematics of UBXD1 constructs used are shown at right.

Dali searches using the VIM-H4 motif did not produce any significant hits, likely due to the structural simplicity of this region. However, examination of other adapters for unannotated structural elements proximal to a VIM suggested that the p97 adapter small VCP/p97-interacting protein (SVIP) might harbor an additional helix in a similar arrangement as in the VIM-H4 of UBXD1^52^. SVIP is a 77-residue minimal adapter that inhibits p97-dependent functions in ERAD, among other functions^52–54^. The SVIP VIM helix is reported to significantly contribute to p97 binding^42^, though a previous study has suggested that additional elements might confer binding affinity^41^. Though no structures of SVIP have been reported, the AlphaFold model of SVIP indeed predicts the VIM helix, as well as an adjacent helix with modest similarity to the UBXD1 H4 (Fig. 5c and Extended Data Fig. 7b). Given the predicted structural similarity to UBXD1 and inhibition of specific cellular functions, we hypothesized that SVIP may similarly remodel p97 D2 contacts and inhibit ATPase activity.

We next purified a previously characterized ASPL construct (ASPL-C, see methods)^46^ and SVIP and analyzed their effect on p97 ATPase activity. ASPL-C potently inhibits p97 ATPase activity, with an IC_50_ of ~97 nM (Fig. 5d). Notably, the complete loss of activity at high ASPL-C concentrations and highly cooperative inhibition (Hill slope ~3) are likely a consequence of hexamer disassembly, as has been previously reported^46^. SVIP also strongly inhibits p97 ATPase activity (IC_50_ ~72 nM). This is striking given its minimal size and indicates that the predicted helix C-terminal to the VIM may contribute to ATPase inhibition through additional interactions with p97. Together these results support a functional conservation between UBXD1, ASPL and SVIP through the inhibition of p97 ATPase activity. Based on our structures and comparison to ASPL and SVIP we postulate that the helical lariat and the H4 helix function as noncanonical control elements that, when paired with well-conserved binding motifs such as UBX and VIM, serve critical functions in ATPase control and p97 remodeling.

To further investigate the helical lariat and H4 in UBXD1, mutations were introduced into the full-length UBXD1 sequence (Extended Data Fig. 7c and methods), and their effect on p97 ATPase activity and structure was determined. We also constructed a double mutant containing both the lariat and H4 changes. Analytical SEC revealed that these constructs bound p97 to a similar extent as did wild type UBXD1 (Extended Data Fig. 7d,e). When tested for p97 ATPase inhibition, the lariat mutant (LX) only modestly increased the IC_50_ compared to wild type UBXD1 (25 nM to 41 nM), indicating its disruption alone does not abolish ATPase inhibition (Fig. 5e). The IC_50_ for the H4 mutant (H4X) was also only modestly shifted compared to wild type (37 nM vs. 25 nM, respectively); rather, the major effect was a ~60% increase in p97 ATPase activity at maximal inhibition (0.42 compared to 0.26 for wild type) (Fig. 5e, dashed lines). This indicates a substantial loss in maximal ATPase inhibition with the H4 mutant. Notably, the double mutant exhibited an increased IC_50_ of 114 nM as well as a similarly elevated maximal inhibition value, indicating a more substantial loss of ATPase inhibition by UBXD1 when both the lariat and H4 are disrupted. Considering the minor IC_50_ effects observed for these mutants individually, these results indicate the lariat and H4 may contribute cooperatively to p97 ATPase control. However, disruption of these elements did not fully abrogate ATPase inhibition, thus additional UBXD1 interactions likely contribute.

Finally, cryo-EM analysis of p97 incubated with ADP and the UBXD1 lariat and H4 mutants was performed to understand structural changes associated with mutation of these motifs (Fig. 6a-d, Extended Data Fig. 7f-j, and Extended Data Table 1). In both datasets, the predominant class contains only VIM UBXD1 density bound in all p97 NTDs, identical to p97:UBXD1^VIM^ in which the p97 hexamer is symmetric and unchanged from the p97^ADP^ state (Extended Data Fig. 7k,l). Additional prevalent classes contain UBXD1 density corresponding to VIM-H4 for the lariat mutant (p97:UBXD1^LX^) and the VIM, PUB, lariat and UBX for the H4 mutant (p97:UBXD1^H4X^, essentially identical to p97:UBXD1^closed^) (Fig. 6a and Extended Data Fig. 7m,n). Density for the mutated regions (lariat or H4) was not observed, indicating loss of these specific interactions was achieved for these variants. Notably, the open-ring state of p97 was not observed in any class, demonstrating that interactions by the lariat and H4 are likely necessary for complete separation of the P1-P6 interface. For p97:UBXD1^LX^ VIM density is better resolved on one protomer (denoted P1) compared to other sites, and following focused classification we identify that this protomer also features H4 density interacting with the D2 domain, as identified in p97:UBXD1^H4^ (Fig. 6b). The clockwise adjacent D2 exhibits strikingly weak density, reminiscent of the D2 flexibility we observed in the p97:UBXD1 closed and open states (Extended Data Fig. 7m). This likely occurs because mutation of Lα3 results in loss of interactions by the UBX and lariat, thereby localizing UBXD1-induced conformational changes to the D2 ring of p97. To explore this state further, 3D variability analysis was performed, revealing a variability mode in which continuous flexibility is observed between two distinct states of the VIM-H4 and adjacent D2 domain. In one state, strong density for both the VIM and H4 is identified and associated with a disordered D2 domain of the clockwise protomer (Fig. 6c and Supplementary Video 5). Conversely, the second state exhibits a well resolved clockwise D2 but weak to no density for the VIM-H4. This analysis indicates these states are in equilibrium and H4 binding correlates with destabilization of the adjacent D2. Thus, these structures further demonstrate that VIM-H4 binding destabilizes p97 through disruption of D2 interprotomer contacts and, together with our ATPase analysis, support a role for the VIM-H4 interaction in D2 hydrolysis control.

**Fig. 6.**
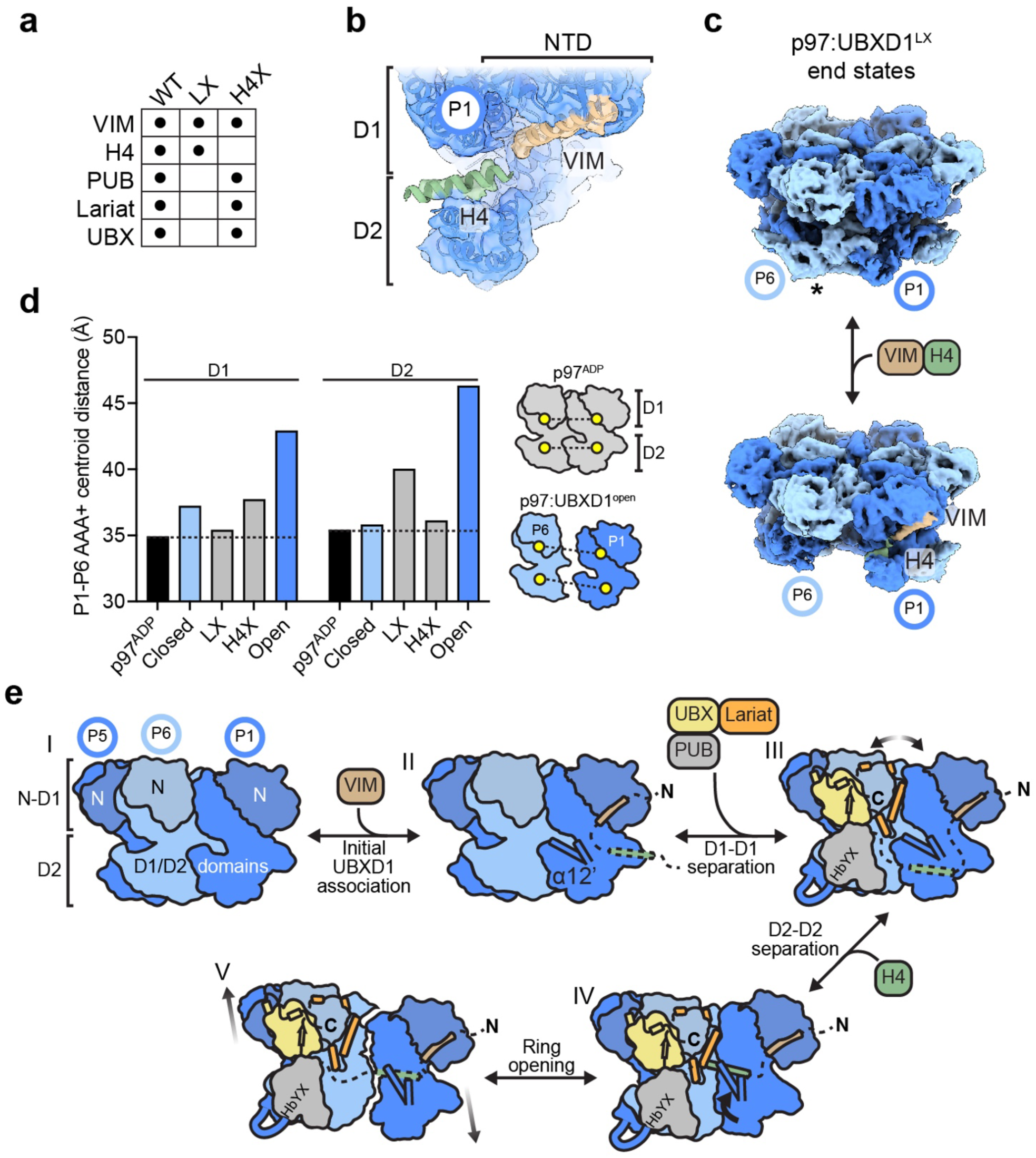
Structural analysis of p97:UBXD1 mutant complexes and model for p97 hexamer remodeling through UBXD1 domain interactions. (**a**) Table of ADP-bound p97:UBXD1 cryo-EM datasets (WT, lariat mutant (LX) and H4 mutant (H4X)) and corresponding UBXD1 domains observed as densities in the reconstructions. (**b**) Unsharpened map and fitted model of the VIM-H4-bound P1 protomer from p97:UBXD1^LX^ (Extended Data Fig. 7f), colored as in Fig. 1. (**c**) First and last frames of the 3D variability analysis output for p97:UBXD1^LX^ showing P6 D2 density (*) but no VIM-H4 in one end state (top) and no P6 D2 in the other when VIM-H4 density is present (bottom). (**d**) P1-P6 interprotomer distances (based on centroid positions) for the D1 and D2 domains of p97^ADP^ (PDB 5FTK), p97:UBXD1^closed^, p97:UBXD1^LX^, p97:UBXD1^H4X^, and p97:UBXD1^open^. Dashed lines represent the minimal distances observed in p97^ADP^. A schematic representing distances calculated is shown (right). (**e**) Model of p97:UBXD1 interactions and structural remodeling of the hexamer. State I: side view of p97^ADP^ (PDB 5FTK), colored as in Fig. 1. NTDs are shaded for clarity. State II: p97:UBXD1^VIM^, in which the VIM initially associates with the NTD of P1. The position of the D2 small subdomain is illustrated by α12’ and an adjacent helix. State III: the p97:UBXD1^closed^ state, in which the UBX, PUB, and helical lariat contact P5 and P6, resulting in the disruption of D1-D1 contacts at the P1-P6 interface. State IV: the p97:UBXD1^H4^ state, in which H4 is positioned on top of the D2 domain of P1, causing it to rotate upward, and displacing a helix from the D2 domain of P6. State V: the p97:UBXD1^open^ state, in which P6 and P1 have completely separated, and all protomers are arranged into a shallow right handed helix.

Considering both p97:UBXD1^LX^ and p97:UBXD1^H4X^ exhibit some P1-P6 asymmetry similar to p97:UBXD1^closed^, we sought to characterize distances between these protomers to further define remodeling of the D1 and D2 by UBXD1. This was achieved by measuring distances between individual AAA+ domains based on centroids calculated from the fitted models (Extended Data Fig. 7o). As expected, the symmetric p97^ADP^ state exhibits the shortest centroid distances (~35Å for D1 and D2) while UBXD1^closed^ and UBXD1^open^ structures show a partial (D1: ~37Å, D2: ~36 Å) and greatly expanded (D1: ~43Å, D2: ~46 Å) separation of the AAA+ domains, respectively (Fig. 6d). As shown above (Fig. 2a), in UBXD1^closed^ the D1 domains are more separated than the D2, which we propose to be caused by lariat binding. Notably, mutation of the lariat (in p97:UBXD1^LX^) decreases the D1-D1 distance relative to p97:UBXD1^closed^ while mutation of H4 (in p97:UBXD1^H4X^) shows no changes in D1-D1, further supporting that the D1 remodeling effects are driven by the lariat interaction. Intriguingly, the D2-D2 distance in p97:UBXD1^LX^ is substantially increased to ~40Å relative to p97:UBXD1^closed^ (at ~36Å). We postulate that this reflects a VIM-H4 interaction that is more pronounced when uncoupled from lariat binding to the D1 (as observed in Fig. 6c), thus supporting that the VIM-H4 interactions indeed contribute to opening the p97 ring through D2 displacement. In sum, these results indicate the helical lariat and H4 independently regulate the D1 and D2 and function as critical p97 hexamer disrupting motifs that are necessary for full UBXD1 remodeling activity.

## DISCUSSION

Adapter proteins of the p97/VCP AAA+ hexamer serve critical roles in binding and regulating function in its many diverse and essential cellular pathways. How adapters may directly regulate p97 structure and mechanism has been an open question. Here we characterized the multi-domain adapter UBXD1 associated with lysosomal and mitochondrial autophagy, among other functions^12,14,17,35,36^. We identify UBXD1 as a potent ATPase inhibitor and determined structures of full-length p97:UBXD1 that reveal how its interactions drive dramatic remodeling and ring-opening of the hexamer. These p97 conformational changes are coordinated by UBXD1 through a network of interprotomer interactions across the N-terminal, D1, and D2 domains. Based on these structures we propose a model describing how UBXD1 interactions coalesce to remodel p97 (Fig. 6e and Supplementary Video 6).

We identify VIM binding to the NTD to be a primary contact given its established interaction^36,39,41,42^ and the prevalence of the p97:UBXD1^VIM^ state (Fig. 6e, state II and Extended Data Fig. 1d), which contains only the VIM helix bound to all protomers. We attribute no p97 structural changes to this interaction alone given this state is unchanged from p97^ADP^. Conversely, interactions made by the helical lariat, including strong contacts by Lα3 along the D1 interprotomer interface, drive remodeling and separation of adjacent D1 domains (Fig. 6e, state III). This is likely supported by UBX binding to the clockwise NTD, given its close connection to the lariat and its conservation with ASPL. Through subclassification of the closed state we identify a class with the UBXD1 H4 helix, which is connected to the VIM by a short linker, bound at the D2 interface (Fig. 4f). This interaction appears critical for separation of the D2 domains given conformational changes identified in this state that destabilize the interface, including displacement of helix α5’ (Fig. 4g). Moreover, variability analysis of the p97:UBXD1^LX^ structure, containing mutations in the lariat, reveals that H4 interactions are dynamic and displace the adjacent D2 (of protomer P6), further supporting a direct role in disrupting the D2 interface (Fig. 6c). Thus, we propose the VIM-H4 specifically functions in p97 D2 remodeling while the combined interactions from the lariat and H4 together drive hexamer opening (Fig. 6e, state V). Additionally, the connecting UBX and VIM domains tether these interactions to the respective NTDs, likely providing additional binding energy to leverage ring opening. Although more flexible, the PUB interaction with the C-terminus of the next clockwise protomer may further support D1-D2 remodeling and ring opening.

Notably, while other UBXD1-bound configurations are identified (Fig. 6d and Extended Data Fig. 1i), the open state is only observed with a singly-bound, wild type UBXD1, thus indicating all UBXD1 contacts across three protomers are required for ring opening. Additional UBXD1 molecules may transiently bind p97, however, opening of the hexamer at one site would likely displace other molecules due to steric interactions and conformational changes across the ring. Given the continuity between the symmetric (p97^ADP^), closed, and open conformations (Supplementary Video 2) and relation to the right-handed spiral adopted by the substrate-bound state^8,9,25^, we propose that a single UBXD1 binds and remodels p97 in this manner to support its substrate processing and translocation cycle. Indeed, UBXD1 cooperates with YOD1 (a deubiquitinase) and PLAA (a ubiquitin-binding adapter) in lysophagy, further supporting its involvement in a substrate-related function^12^. The spiral arrangement facilitated by UBXD1 might allow large substrates or those with complex topologies to enter the central channel for subsequent processing; alternatively, UBXD1 could act as a release factor, allowing stalled substrates to diffuse out of the channel. Furthermore, UBXD1 activity in several pathways appears independent of the ubiquitin-binding adapter UFD1/NPL4^12,14^. This observation, coupled with the unique structural remodeling facilitated by UBXD1, suggests that this adapter enables a function distinct from canonical substrate engagement and processing.

UBXD1’s potent inhibition of p97 ATPase activity is striking and indicates a distinct role for adapters in regulating hydrolysis by p97. Inhibition may result from disruption of D1 and D2 inter-protomer contacts, given the nucleotide pocket resides at the interface and the arginine finger and other intersubunit signaling contacts from the adjacent protomer are lost (Fig. 2d,e)^25^. Indeed, we identify D1 and D2 displacements of 8 and 11 Å, respectively, at the disrupted P6-P1 interface of the open state (Fig. 2d). Additionally, occupancy by the helical lariat of the binding site of UPCDC30245, an allosteric inhibitor that prevents cycling between ADP- and ATP-bound states^19^, indicates that this motif may contribute to ATPase inhibition through an additional mechanism. However, given that the ATPase inhibition activity of the UBXD1 LX mutant is modestly impaired, full inhibition must result through other UBXD1 interactions (Fig. 5e, 6d). Interaction and remodeling of the D2 small subdomain by UBXD1 H4 is also a likely contributor to hydrolysis control and we identify mutation of this helix reduces the maximal inhibition by UBXD1. Hydrolysis inhibition through the D2 is consistent with previous observations that the D2 domain is responsible for the majority of p97 ATPase activity^55^. While we consider the helical lariat and H4 to be primary drivers of ring separation and thus ATPase inhibition, the effects of these remodeling elements are likely buttressed by other UBXD1 domains, given that mutation of the lariat or H4 does not completely ablate UBXD1’s inhibitory activity (Fig. 5e). Therefore, the potent inhibitory effect of UBXD1 may be driven by avidity, without a single interaction domain being specifically responsible for hydrolysis control. Supporting this, many UBXD1 interactions are weak or weakened compared to homologous domains in other adapters. Specifically, H1/H2, though highly conserved (Extended Data Fig. 8), weakly interacts by NMR^36–38,56^, its VIM is lacking an Arg residue present in many other adapters^41,42^, its UBX has residues in the S3/S4 loop mutated and does not bind the NTD in isolation^17,39,43^, and H4 also weakly binds as measured by NMR^36^. Notably, UBXD1-mediated inhibition of ATPase activity does not necessitate an overall inhibitory role of this adapter; UBXD1 could potentially stabilize a stalled, ATPase-inhibited state until association of a substrate, at which point hydrolysis-dependent substrate processing might occur.

We propose UBXD1, ASPL, and SVIP are structurally related adapters with conserved motifs that mediate distinct effects on p97 activity. In addition to being potent inhibitors of p97 ATPase activity, these adapters appear to not bind substrate or possess enzymatic activity, suggesting their primary activity is modulation of p97 structure rather than direct involvement in substrate engagement. In contrast to many other AAA+ proteins, p97 requires an extensive set of adapter proteins, including those that deliver substrates, to facilitate its functions. This is likely due to the relative stability of the p97 hexamer, which adopts a stable, planar conformation in the absence of substrate and even nucleotide^18,57^. It is therefore reasonable to conclude that p97 relies on a set of adapters to structurally remodel the hexamer for various purposes, in the same manner as its dependence on adapters directly involved in substrate processing. We identify the helical lariat and H4-like sequences to be critical control elements in these adapters. The different degrees of remodeling of lariat-containing adapters (complete hexamer disassembly with ASPL compared to intact hexamers with UBXD1) likely reflect the different assembles of p97-interacting domains in these two adapters. In addition to the lariat-UBX module, ASPL has a UBXL domain and a SHP box that both bind the NTD, while UBXD1 features a much larger complement of domains that flank the lariat on both sides. These extra domains could potentially ‘hold’ the hexamer together during binding, preventing complete dissociation, possibly to facilitate an as-yet unknown step of substrate processing. Indeed, a recent study revealed that ASPL-mediated hexamer disassembly enables binding and modification by the methyltransferase VCPKMT, suggesting that disruptions of hexamer architecture are biologically relevant^58^. Likewise, the extra p97-interacting domains of UBXD1 compared to SVIP indicate a more sophisticated function than merely ATPase inhibition, as SVIP similarly inhibits hydrolysis with only VIM and H4-like sequences. In sum, the characterization of adapters as structural modulators of p97 reported here and the large number of still-uncharacterized p97-interacting proteins suggest that there are more classes of adapters with distinct effects on p97 activity yet to be discovered, likely with novel regulatory effects on unfoldase function.

## ACKNOWLEDGMENTS

This work was supported by NIH grants F31GM142279 (to J.R.B.), R01GM130145 (to M.R.A.), and R01GM138690 (to D.R.S.).

## AUTHOR CONTRIBUTIONS

J.R.B. expressed and purified proteins, performed biochemical and cryo-EM experiments, built models, developed figures, and wrote and edited the manuscript. C.R.A. expressed and purified proteins, performed biochemical experiments, and edited the manuscript. M.R.T. expressed and purified proteins and performed cryo-EM experiments. E.T. operated electron microscopes and assisted with data collection. A.T. expressed proteins. M.R.A. designed and supervised biochemistry and edited the manuscript. D.R.S. designed and supervised the project and wrote and edited the manuscript.

## RESOURCE AVAILIBITY

### Materials availability

Requests for resources and reagents should be directed to Daniel R. Southworth.

### Data availability

Cryo-EM densities have been deposited at the Electron Microscopy Data Bank under accession codes EMD: 28982 (p97:UBXD1^closed^), EMD: 28983 (p97:UBXD1^open^ composite), EMD: 28984 (p97:UBXD1^open^ consensus), EMD: 28985 (p97:UBXD1^open^ P1 focused map), EMD: 28986 (p97:UBXD1^open^ P6 focused map), EMD: 28987 (p97:UBXD1^VIM^), EMD: 28988 (p97:UBXD1^meta^), EMD: 28989 (p97:UBXD1^para^), EMD: 28990 (p97:UBXD1-PUB_in_), EMD: 28991 (p97:UBXD1^H4^), and EMD: 28992 (p97:UBXD1^LX^). Atomic coordinates have been deposited at the Protein Data Bank under accession codes PDB: 8FCL (p97:UBXD1^closed^), PDB: 8FCM (p97:UBXD1^open^), PDB: 8FCN (p97:UBXD1^VIM^), PDB: 8FCO (p97:UBXD1^meta^), PDB: 8FCP (p97:UBXD1^para^), PDB: 8FCQ (p97:UBXD1-PUB_in_), PDB: 8FCR (p97:UBXD1^H4^), and PDB: 8FCT (p97:UBXD1^LX^).

## METHOD DETAILS

### Molecular cloning

The coding sequence of full-length human UBXD1 was cloned with an N-terminal 6xHis tag, MBP tag, and TEV protease cleavage site into an insect cell expression vector (Addgene plasmid #55218). The same expression construct was cloned into a bacterial expression vector (Addgene plasmid #29708). ASPL-C (residues 313-553) and full-length SVIP were cloned into the same expression vector. The NEB Q5 Site-Directed Mutagenesis kit was used to introduce mutations into the UBXD1 construct. The UBXD1 lariat was mutated by making four charge reversals in Lα3 predicted to disrupt contacts with P1 and P6 (E299R/R302E/R307E/E312R), given this helix makes the most significant contacts with p97. As we could not obtain high-resolution structural information about H4, we scrambled the sequence of this helix rather than making point mutants, using Peptide Nexus Sequence Scrambler (https://peptidenexus.com/article/sequence-scrambler). This resulted in the sequence 75-QSRDVTQERIQNKAVLTEA-93.

### Protein expression and purification

p97 was expressed and purified as described previously^55^. Briefly, BL21-Gold(DE3) chemically competent *E. coli* (Agilent) were transformed with pET15b p97, encoding full-length p97 with an N-terminal 6xHis tag, grown in 2xYT media supplemented with 100 µg/mL carbenicillin, and induced with 0.5 mM isopropyl β-D-1-thiogalactopyranoside (IPTG) at 20°C overnight. Cells were harvested and lysed by sonication in lysis buffer (50 mM Tris pH 8.0, 250 mM NaCl, 10 mM imidazole, 0.5 mM Tris(2-carboxyethyl)phosphine (TCEP), 1 mM phenylmethylsulfonyl fluoride (PMSF)) supplemented with cOmplete Protease Inhibitor Cocktail, EDTA-free (Roche) then clarified by centrifugation. The supernatant was then incubated with HisPur Ni-NTA resin (Thermo Scientific), and p97 was eluted with nickel elution buffer (lysis buffer supplemented with 320 mM imidazole, no PMSF). The eluate was supplemented with TEV protease and dialyzed overnight at 4°C into p97 dialysis buffer (10 mM Tris pH 8.0, 100 mM NaCl, 1 mM dithiothreitol (DTT)). The following day, the cleavage product was passed through fresh Ni-NTA resin, and the flowthrough was concentrated and applied to a HiLoad 16/600 Superdex 200 pg size exclusion chromatography (SEC) column (GE Healthcare) equilibrated in p97 SEC buffer (25 mM HEPES pH 7.4, 150 mM KCl, 5 mM MgCl_2_, 0.5 mM TCEP). Fractions containing p97 were concentrated to >200 µM, filtered, and flash frozen in liquid nitrogen.

Initial attempts at expression of UBXD1 in *E. coli* resulted in large amounts of insoluble material. Therefore, to obtain amounts sufficient for initial studies, a UBXD1 construct with a TEV-cleavable N-terminal 6xHis-MBP tag was expressed in Sf9 insect cells using standard methods. This protocol (including cleavage of the 6xHis-MBP tag) yielded sufficient material for preliminary cryo-EM studies, and was used for the p97/UBXD1^WT^/ADP dataset. Thereafter, an optimization campaign for soluble expression of 6xHis-MBP-UBXD1 in *E. coli* was performed, which resulted in the following protocol. *E. coli*-derived UBXD1 and mutants thereof were used for all biochemical experiments, and the p97/UBXD1/ATPγS, p97/UBXD1^LX^/ADP and p97/UBXD1^H4X^/ADP datasets. BL21-Gold(DE3) chemically competent *E. coli* (Agilent) were transformed with the UBXD1 expression vectors and used for large-scale expression. Cells were grown in 2xYT media supplemented with 100 µg/mL carbenicillin at 37°C until OD_600_ reached ~1.25, then protein expression was induced with 0.5 mM IPTG, and grown for 1 hour at 37°C. Cells were rapidly cooled in an ice bath for 10 min, then harvested by centrifugation at 10,000 RCF and stored at −80°C until use. All subsequent steps were performed at 4°C. Pellets were resuspended in lysis buffer (see above) supplemented with cOmplete Protease Inhibitor Cocktail, EDTA-free (Roche), and lysed by sonication. Lysates were clarified by centrifugation at 85,000 RCF and incubated with HisPur Ni-NTA resin (Thermo Scientific) for 15 min. The resin was washed with nickel wash buffer (lysis buffer without imidazole or PMSF) and eluted with nickel elution buffer (see above). The eluate was concentrated, filtered, and applied to a HiLoad 16/600 Superdex 200 pg SEC column (GE Healthcare) equilibrated in adapter SEC buffer (25 mM HEPES pH 7.4, 150 mM KCl, 5% glycerol (v/v), 0.5 mM TCEP). TEV protease was added to fractions containing MBP-UBXD1 and incubated overnight without agitation. The following day, the sample was passed through a 5 mL MBPTrap HP column (GE Healthcare) to remove the cleaved 6xHis-MBP tag, and the flowthrough was concentrated before 15x dilution with anion exchange (AEX) binding buffer (25 mM HEPES pH 7.5, 0.5 mM TCEP). The diluted sample was then applied to a 1 mL HiTrap Q HP column (GE Healthcare), and UBXD1 was eluted with a 0-50% gradient of AEX elution buffer (AEX binding buffer supplemented with 1000 mM KCl), concentrated to >200 µM, and flash frozen in liquid nitrogen.

To express ASPL-C and SVIP, BL21-Gold(DE3) were transformed with pET MBP-ASPL-C and pET MBP-SVIP, and grown in Terrific Broth supplemented with 100 µg/mL ampicillin at 37°C until OD_600_ reached ~2, then protein expression was induced with 0.4 mM IPTG, and grown overnight at 18°C. Cells were harvested by centrifugation at 4,000 RCF and stored at −80°C or processed immediately. All subsequent steps were performed at 4°C. Cell lysis and nickel immobilized metal affinity chromatography were performed as for UBXD1. TEV protease was added to the eluates, and the solutions were dialyzed overnight in adapter dialysis buffer (25 mM Tris pH 8.0, 150 mM NaCl, 0.5 mM TCEP). The following day, the samples were passed through fresh Ni-NTA resin to remove the 6xHis-MBP tags, concentrated, and applied to a HiLoad 16/600 Superdex 200 pg SEC column equilibrated in adapter SEC buffer. Fractions containing adapter proteins were concentrated to >200 µM and flash frozen in liquid nitrogen.

Purity of all proteins was verified by SDS-PAGE and concentration was determined using the Pierce BCA Protein Assay Kit (Thermo Scientific).

### ATPase assays

The ATPase assay protocol was modified from previously published methods^60^. In an untreated 384-well microplate (Grenier 781101), 50 µL solutions were prepared to a final concentration of 10 nM p97 hexamer, variable adapter (UBXD1 and mutants, ASPL-C, and SVIP), and 200 µM ATP (Thermo Fisher Scientific) in ATPase buffer (25 mM HEPES pH 7.4, 100 mM KCl, 3 mM MgCl_2_, 1 mM TCEP, 0.1 mg/mL BSA). ATP was added last to initiate the reaction, and the solutions were incubated at room temperature until 8% substrate hydrolysis was achieved. To quench the reaction, 50 µL of BIOMOL Green (Enzo Life Sciences) was added and allowed to develop at room temperature for 25 min before reading at 620 nm.

### Analytical size exclusion chromatography

60 µL samples (10 µM p97 monomer, 20 µM UBXD1, and 5 mM nucleotide where applicable) were prepared in p97 SEC buffer and incubated on ice for 10 minutes. Samples were filtered and injected on a Superose 6 Increase 3.2/300 column (GE Healthcare) equilibrated in p97 SEC buffer and operated at 8°C. 100 µL fractions were collected and analyzed by SDS-PAGE with Coomassie Brilliant Blue R-250 staining (Bio-Rad).

### Cryo-EM data collection and processing

For all p97/UBXD1 datasets, 10 µM p97 monomer and 20 µM UBXD1 were incubated with 5 mM ADP or ATPγS in p97 SEC buffer for 10 minutes on ice before vitrification. A 3 µL drop was applied to a glow-discharged (PELCO easiGlow, 15 mA, 2 min) holey carbon grid (Quantifoil R1.2/1.3 on gold 200 mesh support), blotted for 3-4 seconds with Whatman Grade 595 filter paper (GE Healthcare), and plunge frozen into liquid ethane cooled by liquid nitrogen using a Vitrobot (Thermo Fisher Scientific) operated at 4°C and 100% humidity. Samples were imaged on a Titan Krios TEM (Thermo Fisher Scientific) operated at 300 kV and equipped with a BioQuantum K3 Imaging Filter (Gatan) using a 20 eV zero loss energy slit. Movies were acquired with SerialEM^61^ in super-resolution (UBXD1^WT^/ADP, UBXD1^H4X^/ADP, UBXD1^WT^/ATPγS) or counted (UBXD1^LX^/ADP) mode at a calibrated magnification of 59,952x, corresponding to a physical pixel size of 0.834 Å. A nominal defocus range of −0.8 to −1.8 µm was used with a total exposure time of 2 sec fractionated into 0.255 sec frames for a total dose of 43 e^−^/Å^2^ at a dose rate of 15 e^−^/pix/s. Movies were subsequently corrected for drift and dose-weighted using MotionCor2^62^, and the micrographs collected in super-resolution mode were Fourier cropped by a factor of 2.

For the p97/UBXD1^WT^/ADP sample, a total of 22,536 micrographs were collected and initially processed in cryoSPARC^63^. After Patch CTF estimation, micrographs were manually curated to exclude those of poor quality, followed by blob-based particle picking, 2D classification, *ab initio* modeling, and initial 3D classification. Three classes of interest from the initial 3D classification were identified, corresponding to a closed-like state (class 1), an open-like state (class 2), and p97:UBXD1^VIM^ (class 3). For p97:UBXD1^VIM^, 100,000 particles were randomly selected and refined with C6 symmetry imposed. For the open state, refinement of all particles produced a map with poor resolution for protomers P1 and P6, so focused classification without image alignment (skip-align) of these individual protomers was performed in RELION^64^, followed by masked local refinement in cryoSPARC. A composite map of these two local refinements and protomers P2-P5 from the consensus map was generated in UCSF Chimera by docking the local refinement maps into the consensus map, zoning each map by a radius of 4 Å using the associated chains, and summing the aligned volumes^65^. For the closed state, refinement of the closed-like particles revealed additional UBXD1 density at protomers other than P1 and P6, so skip-align focused classification using a mask encompassing protomers with additional density was performed in RELION, followed by homogenous refinement in cryoSPARC, yielding the p97:UBXD1^meta^ and p97:UBXD1^para^ states. The class from focused classification corresponding to a singly-bound hexamer was also refined in cryoSPARC, and then subjected to skip-align focused classification of P1 and P6 in RELION, followed by homogenous refinement in cryoSPARC. This yielded the p97:UBXD1^closed^ and p97:UBXD1^H4^ states. To obtain better density for the PUB domain, all particles from the closed-like class from the initial 3D classification were used for skip-align focused classification of the PUB and UBX domain in RELION, followed by homogenous refinement in cryoSPARC. This yielded p97:UBXD1-PUB_out_ and p97:UBXD1-PUB_in_. To visualize variability in this dataset, 3D Variability Analysis in cryoSPARC was performed using particles from the closed-like and open-like states.

For the p97/UBXD1^WT^/ATPγS dataset, a total of 9,498 micrographs were collected and initially processed as for p97/UBXD1^WT^/ADP. An initial 3D classification revealed three classes of interest: a state resembling p97:UBXD1^closed^, a state resembling a fully ATPγS-bound p97 hexamer with NTDs in the up state, and a state resembling the p97:UBXD1^VIM^ state. Homogenous refinement with defocus refinement was performed for each of these classes, and the resolution in all maps was sufficient to assign nucleotide density as ATPγS in all nucleotide pockets in all protomers in all structures.

For the p97/UBXD1^LX^/ADP sample, a total of 6,330 micrographs were collected and initially processed as for p97/UBXD1^WT^/ADP. An initial 3D classification revealed two high-resolution classes, one featuring density for the UBXD1 VIM in all NTDs (class 1), and the other featuring stronger density for the VIM in one NTD than in the others (class 2). In this class the D2 domain of the protomer counterclockwise from the best VIM-bound protomer had significantly weaker density, indicating flexibility. Homogenous refinement of class 1 produced a map essentially identical to p97:UBXD1^VIM^. Homogenous refinement of class 2 produced a map with density weak density putatively corresponding to H4 on the strong VIM-bound protomer, so skip-align focused classification of the D2 domain of this protomer was performed in RELION, followed by a final homogenous refinement in cryoSPARC of the best class. This yielded a map with improved VIM and H4 density. To visualize variability in this dataset, 3D Variability Analysis in cryoSPARC was performed using particles from the class 2 refinement (pre-focused classification).

For the p97/UBXD1^H4X^/ADP sample, a total of 12,418 micrographs were collected and initially processed as for p97/UBXD1^WT^/ADP. An initial 3D classification revealed three high-resolution classes, one featuring density for the UBXD1 VIM in all NTDs (class 1), and the other two (classes 2 and 3) featuring stronger density for the VIM and additional UBXD1 domains, including the PUB, helical lariat, and UBX. These classes strongly resemble p97:UBXD1^closed^. Homogenous refinement of class 1 produced a map essentially identical to p97:UBXD1^VIM^. Homogenous refinement of classes 2 and 3 combined produced a map with density for the VIM, PUB, lariat, and UBX, but without H4 density or associated movements of the D2 domains. To identify particles with H4 density, skip-align focused classification of protomers P1 and P6 was performed in RELION, which did not reveal any classes with density attributable to H4.

### Molecular modeling

To generate the model for p97:UBXD1^closed^, a model of the p97^ADP^ hexamer^19^ and the AlphaFold model of UBXD1^44,45^ were docked into the map using UCSF Chimera^65^ and ISOLDE^66^ in UCSF ChimeraX^67^, followed by refinement using Rosetta Fast Torsion Relax. Models for all other structures were generated by docking individual chains from the closed model, followed by refinement using Rosetta Fast Torsion Relax. Coot^68^, ISOLDE, and Phenix^69^ were used to finalize all models. Sidechains for H4 residues in p97:UBXD1^H4^ and p97:UBXD1^LX^ are omitted due to low resolution.

### UBXD1 sequence alignment

UBXD1 protein sequences were aligned in MUSCLE and the resulting alignment was visualized in MView^70^.

### Data analysis and figure preparation

Biochemical data was analyzed and plotted using Prism 9.3.1 (GraphPad). Figures were prepared using Adobe Illustrator, UCSF Chimera, and UCSF ChimeraX^65,67^.

## EXTENDED DATA

**Extended Data Fig. 1.**
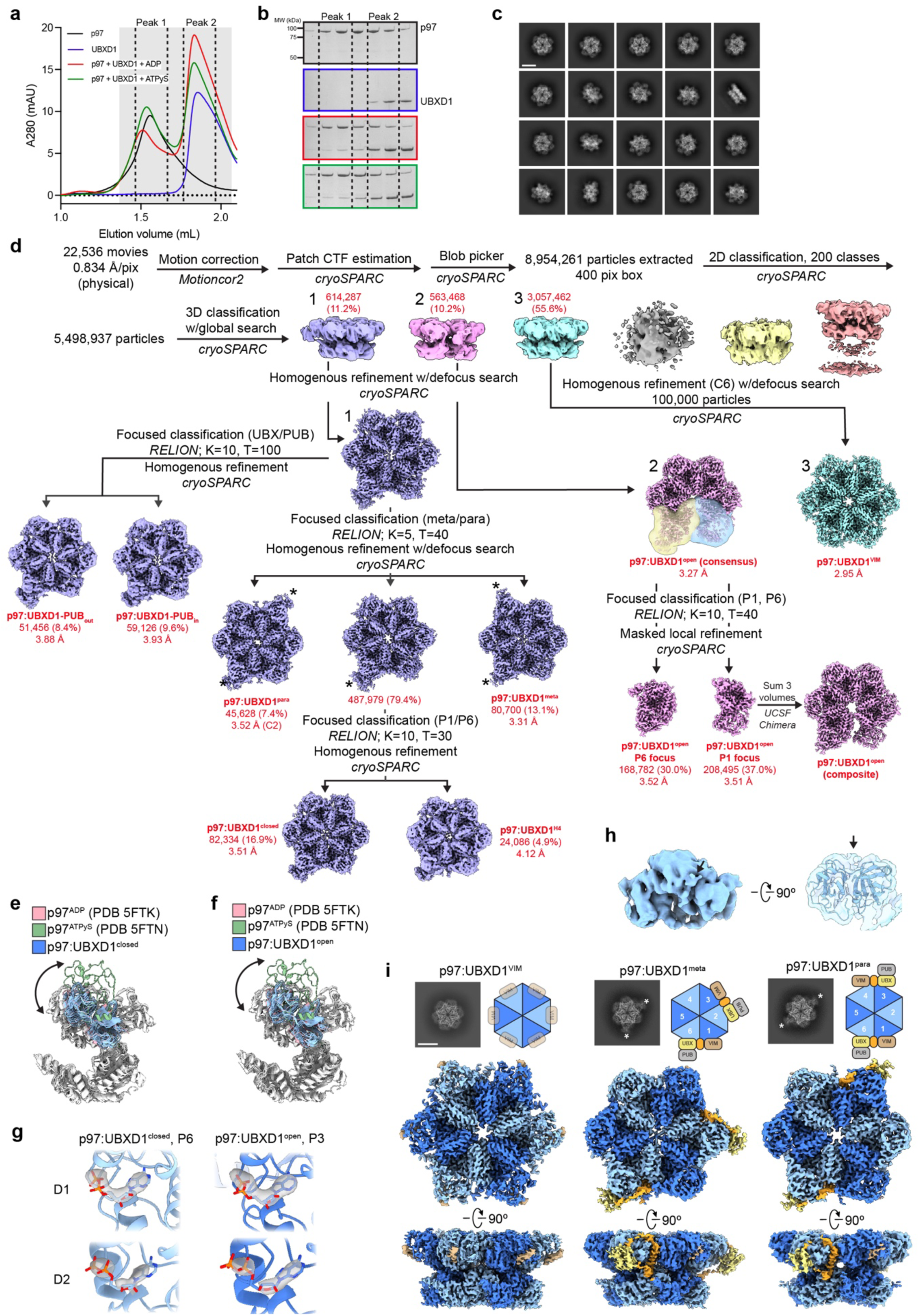
Biochemical and cryo-EM analysis of the p97:UBXD1 interaction. (**a**) SEC traces of p97:UBXD1 complexes, showing a left shift in peak elution volume for p97 samples with UBXD1. Fractions in the shaded range were analyzed by SDS-PAGE. No p97 monomer peak was observed with UBXD1 incubation. (**b**) Coomassie Brilliant Blue-stained SDS-PAGE gels of fractions from SEC runs in (**a**). (**c**) Representative 2D class averages of the p97/UBXD1^WT^/ADP dataset (scale bar equals 100 Å). No p97 monomers were identified during 2D classification. (**d**) Processing workflow for structures obtained from the p97/UBXD1^WT^/ADP dataset. Masks used for the P1 and P6 focused classification and masked local refinement of p97:UBXD1^open^ are shown in transparent blue and yellow, respectively. (**e**) Overlay of all protomers from p97:UBXD1^closed^ (blue) with a protomer in the ADP-bound, down NTD conformation (pink, PDB 5FTK) and a protomer in the ATPγS-bound, up NTD conformation (green, PDB 5FTN), aligned by the D1 large subdomain (residues 211-368). For all protomers, the NTDs are colored, and the D1 and D2 domains are white. (**f**) As in (**e**), but depicting protomers from p97:UBXD1^open^ (blue). (**g**) Nucleotide densities for representative D1 and D2 pockets in p97:UBXD1^closed^ and p97:UBXD1^open^. (**h**) Representative additional density in NTD corresponding to a VIM helix (unsharpened map of P4 in p97:UBXD1^open^). (**i**) Cartoons, top view projections of sharpened maps showing UBX/PUB density (*), and sharpened maps of p97:UBXD1^VIM^, p97:UBXD1^meta^, and p97:UBXD1^para^ (scale bar equals 100 Å). In p97:UBXD1^VIM^, the VIM density is depicted as a difference map of p97:UBXD1^VIM^ and a map generated from a model without VIM helices.

**Extended Data Fig. 2.**
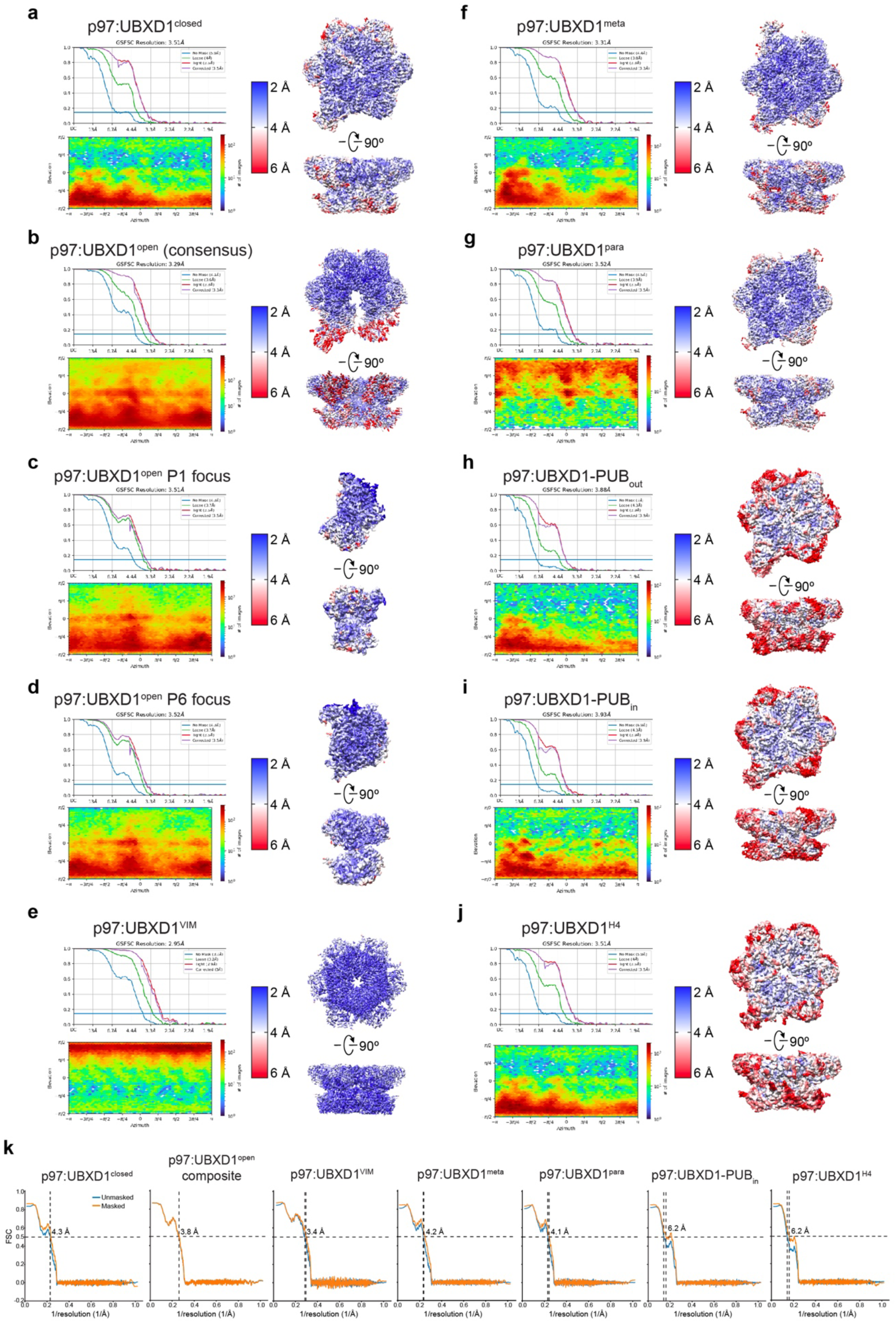
Cryo-EM densities and resolution estimation from the ADP-bound p97:UBXD1^WT^ dataset. (**a** to **j**) Fourier shell correlation (FSC) curves, particle orientation distribution plots, and sharpened density maps colored by local resolution (0.143 cutoff) for (**a**) p97:UBXD1^closed^, (**b**) p97:UBXD1^open^ (consensus map), (**c**) p97:UBXD1^open^ P1 focus, (**d**) p97:UBXD1^open^ P6 focus, (**e**) p97:UBXD1^VIM^, (**f**) p97:UBXD1^meta^, (**g**) p97:UBXD1^para^, (**h**) p97:UBXD1-PUB_out_, (**i**) p97:UBXD1-PUB_in_, and (**j**) p97:UBXD1^H4^. (**k**) Map-model FSC curves for p97:UBXD1^closed^, p97:UBXD1^open^ (composite map), p97:UBXD1^VIM^, p97:UBXD1^meta^, p97:UBXD1^para^, p97:UBXD1-PUB_in_, and p97:UBXD1^H4^. Displayed model resolutions were determined using the masked maps.

**Extended Data Fig. 3.**
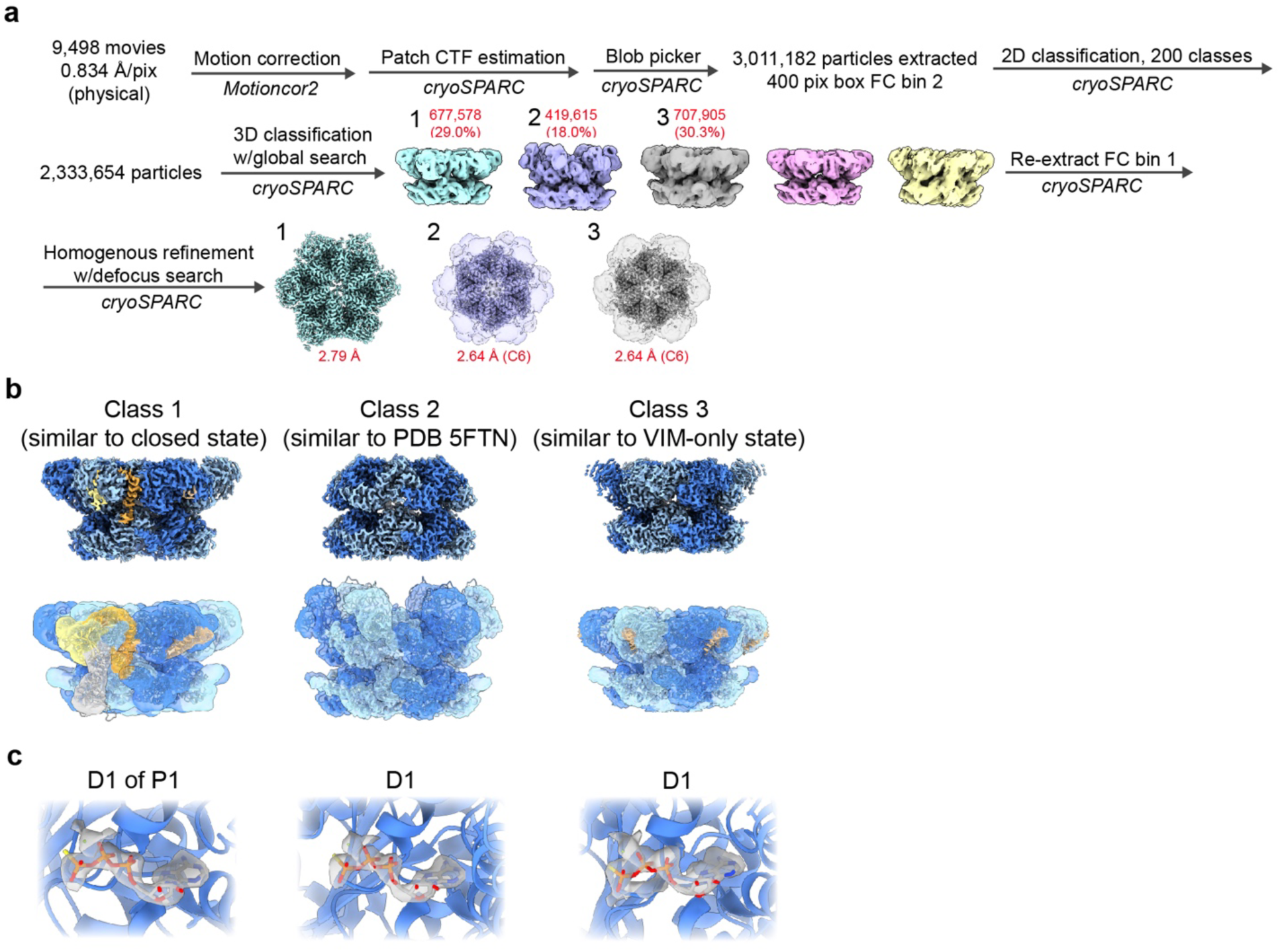
Cryo-EM analysis of p97:UBXD1 incubated with ATPγS. (**a**) Processing workflow for structures obtained from the p97/UBXD1^WT^/ATPγS dataset. (**b**) (Top row) sharpened maps of class 1-3 refinements. (Bottom row) p97:UBXD1^closed^ model overlaid with filtered map, unsharpened map, ATPγS-bound p97 hexamer with NTDs in the up state (PDB 5FTN) overlaid with the class 2 unsharpened map, and p97:UBXD1^VIM^ model overlaid with the class 3 unsharpened map, respectively. All maps are colored as in Fig. 1. (**c**) Representative nucleotide densities for class 1-3 refinements (sharpened maps), showing clear γ-phosphate and Mg^2+^ density. The nucleotide and binding pocket from PDB 5FTN are shown for clarity.

**Extended Data Fig. 4.**
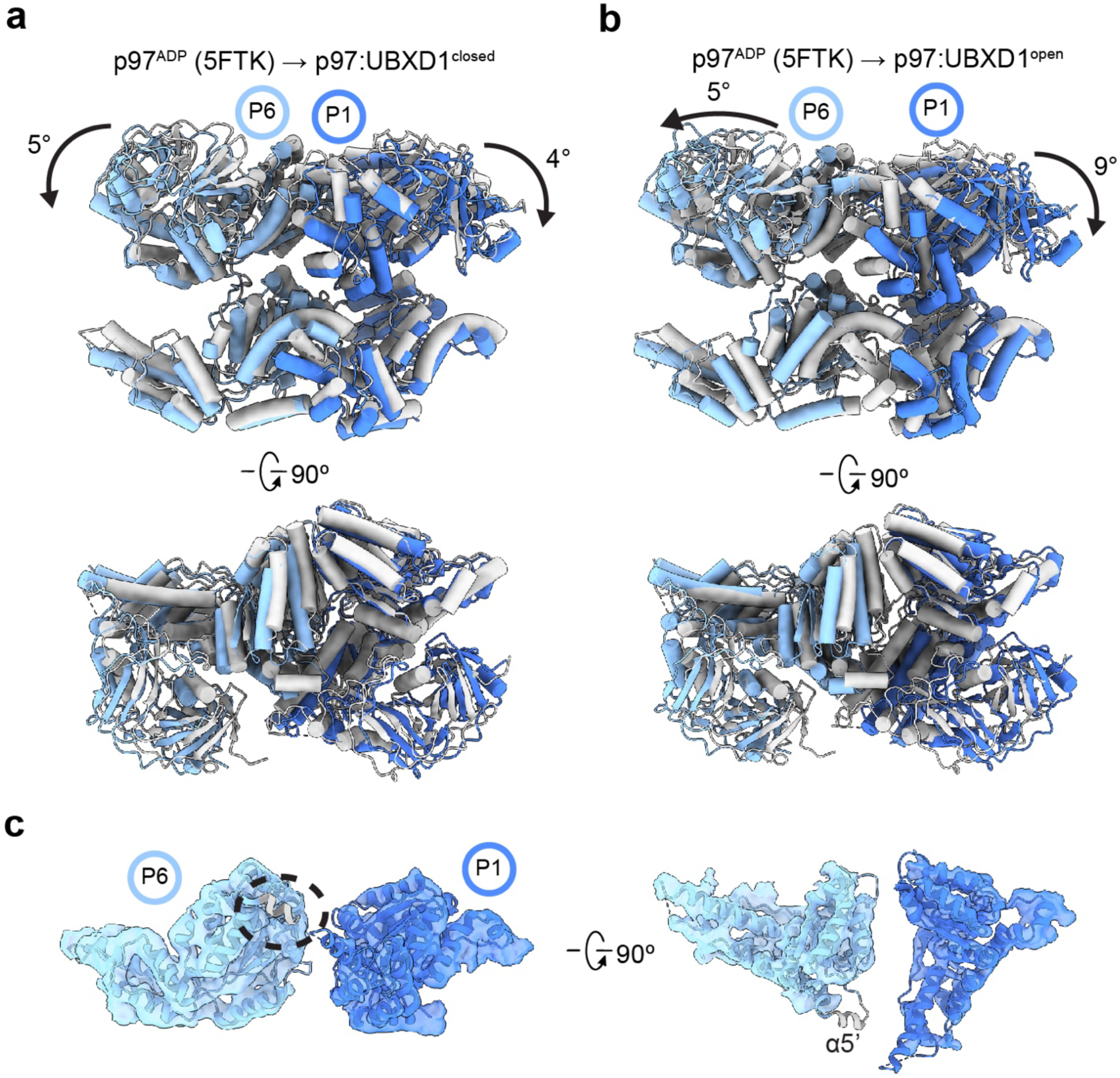
Changes in p97:UBXD1^closed^ and p97:UBXD1^open^. (**a**) Overlay of protomers P1 (dark blue) and P6 (light blue) from p97:UBXD1^closed^, aligned to protomers P3 and P4 from PDB 5FTK. P1 and P6 protomers from 5FTK are shown in gray. (**b**) As in (**a**), but depicting p97:UBXD1^open^ protomers. (**c**) Unsharpened map of the D2 domains of protomers P1 and P6 of p97:UBXD1^open^, overlaid with the D2 domain from p97^ADP^ on P6, showing lack of density for helix α5’ (gray, encircled) normally contacting the counterclockwise D2 domain. The D2 domain of P1 of the open state is shown for clarity.

**Extended Data Fig. 5.**
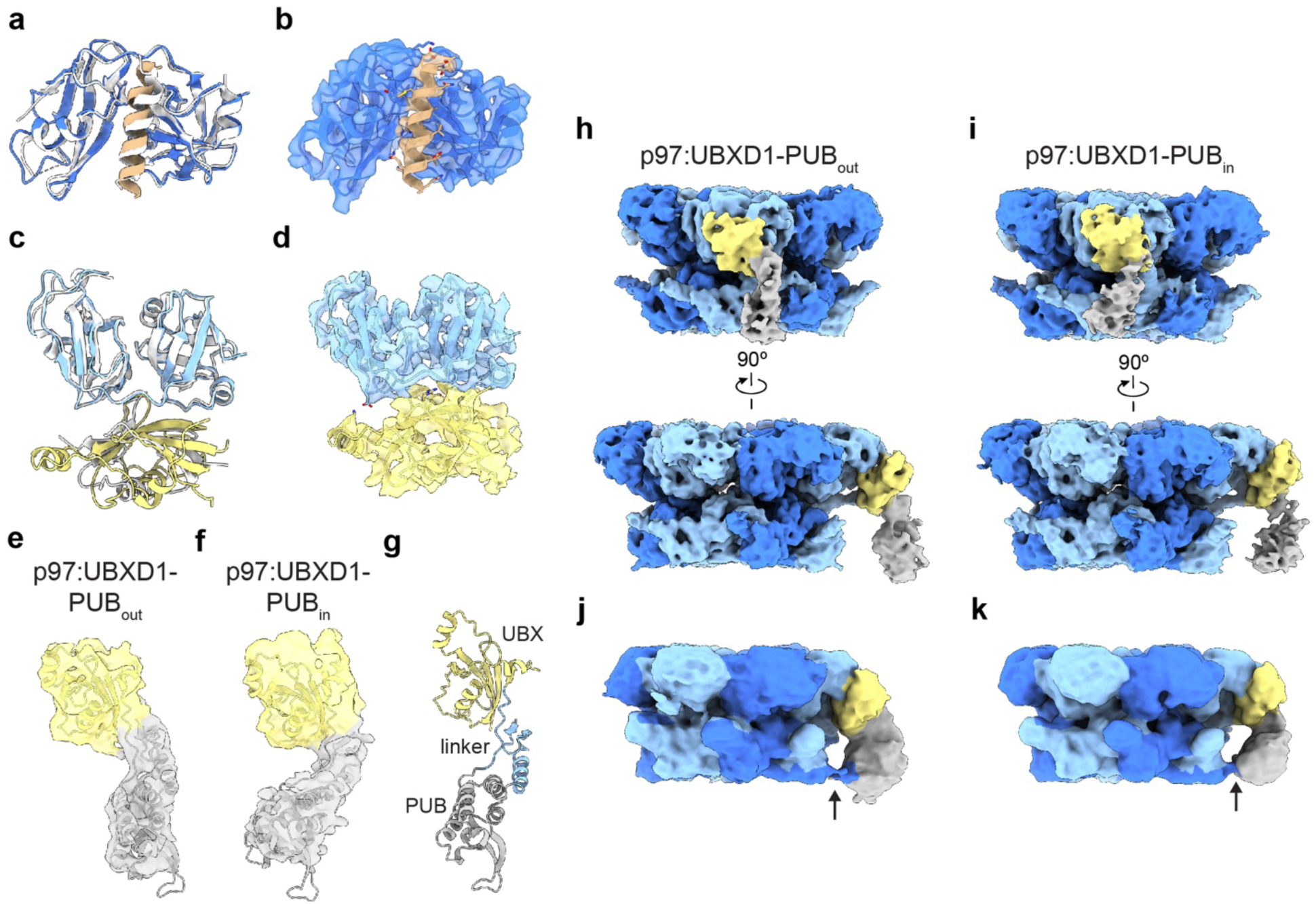
VIM, UBX, and PUB comparisons and validation. (**a**) Overlay of NTD-VIM from p97:UBXD1^closed^ (colored) and gp78 (PDB 3TIW, white). (**b**) Map and model of the NTD and VIM from p97:UBXD1^closed^, colored as in (**a**). (**c**) Overlay of NTD-UBX from p97:UBXD1^closed^ (colored) and UBXD7 (PDB 5X4L, white). (**d**) Map and model of the NTD and UBX from p97:UBXD1^closed^, colored as in (**c**). (**e**) Unsharpened, zoned map and model of UBX and PUB from p97:UBXD1-PUB_out_. (**f**) Unsharpened, zoned map and model of UBX and PUB from p97:UBXD1-PUB_in_. (**g**) Model of the UBX (yellow), PUB (gray), and UBX-PUB linker (light blue) from p97:UBXD1^closed^. (**h**) Unsharpened map of p97:UBXD1-PUB_out_.The VIM and helical lariat are colored the same as their corresponding protomers for clarity. (**i**) Unsharpened map of p97:UBXD1-PUB_in_. The VIM and helical lariat are colored the same as their corresponding protomers for clarity. (**j**) Filtered map of p97:UBXD1-PUB_out_, colored as in (**d**), showing weak density connecting the PUB and P5 CT tail. (**k**) As in (**j**), but for p97:UBXD1-PUB_in_.

**Extended Data Fig. 6.**
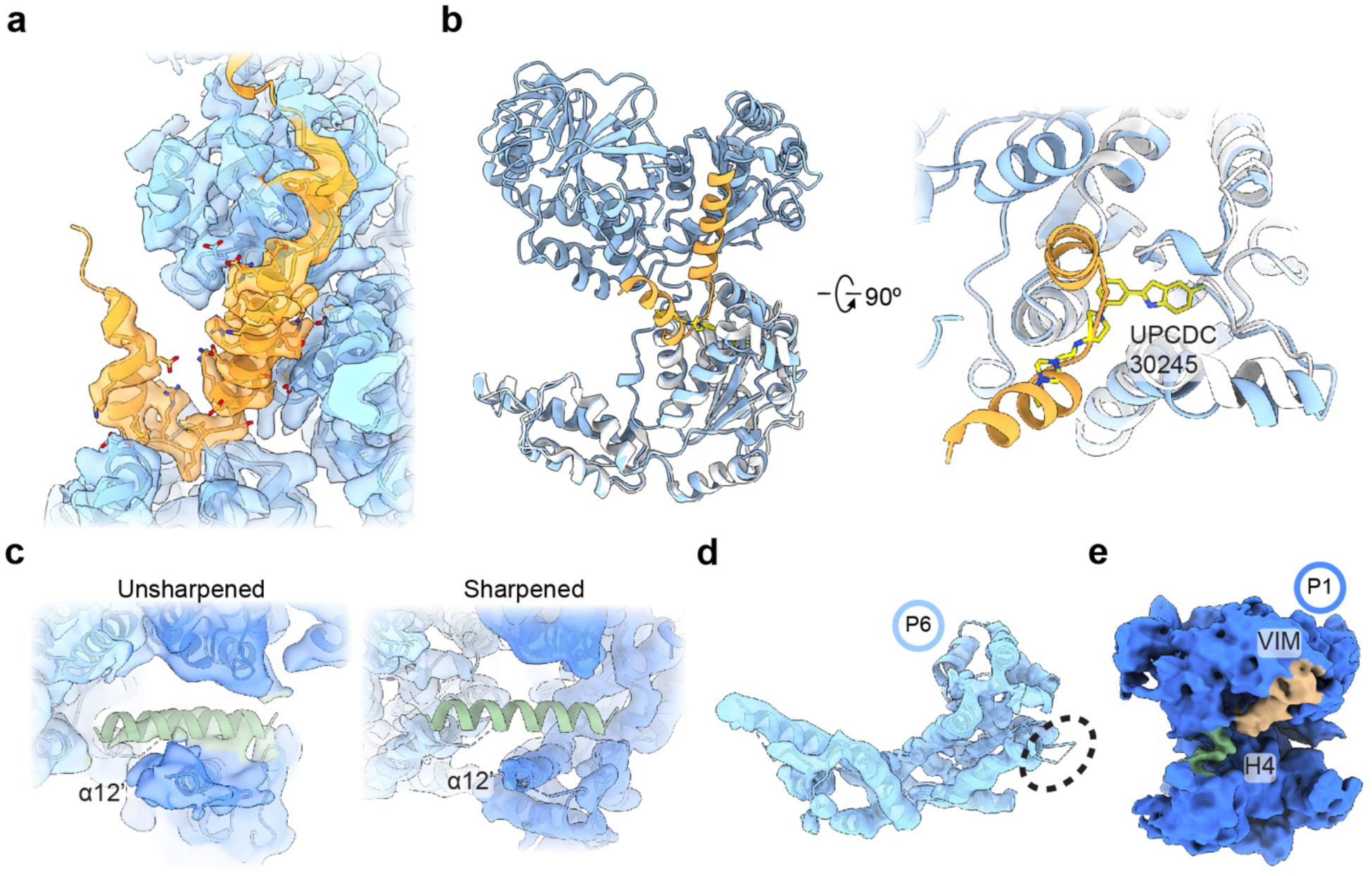
Validation of the helical lariat, UPCDC30245 binding, and additional structural features of p97:UBXD1^H4^ and p97:UBXD1^open^. (**a**) Sharpened map and model of Lα2-4, connecting strands of the helical lariat, and adjacent regions of P6 of p97:UBXD1^meta^, colored as in Fig. 1. (**b**) Overlay of P6 (blue) and Lα3-4 (orange) from p97:UBXD1^closed^ with a p97 protomer (white) bound to UPCDC30245 (yellow) (PDB 5FTJ), aligned by the D2 domain (residues 483-763). (**c**) View of H4 and surrounding p97 density in p97:UBXD1^H4^ (left: unsharpened, right: sharpened) overlaid with the model for this state. (**d**) Sharpened map and model of the D2 domain of protomer P6 of p97:UBXD1^H4^, showing lack of density for α5’ (encircled). (**e**) Unsharpened map of protomer P1 of p97:UBXD1^open^. Density putatively corresponding to H4 is colored in green.

**Extended Data Fig. 7.**
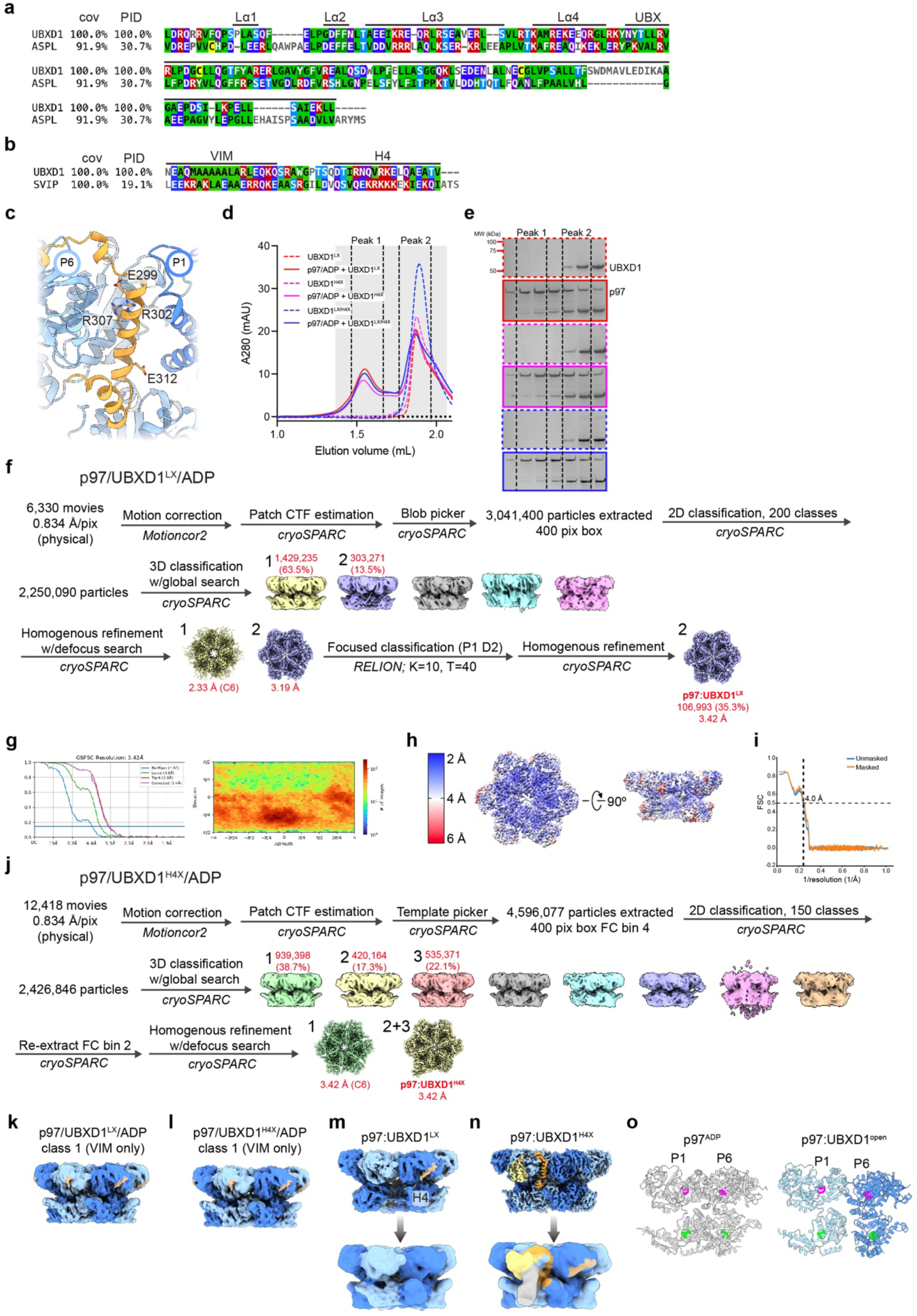
Conservation and structural analysis of UBXD1, ASPL, and SVIP. (**a**) Alignment of UBX-lariat sequences from UBXD1 (residues 270-441) and ASPL (residues 318-495). Structural elements in the UBXD1 sequence are indicated above. Cov = covariace relative to the human sequence, Pid = percent identity relative to the human sequence. (**b**) Alignment of VIM-H4 sequences from UBXD1 (residues 50-93) and SVIP (residues 18-64). Structural elements in the UBXD1 sequence are indicated above. (**c**) Residues mutated in Lα3 of the UBXD1 helical lariat, shown on p97:UBXD1^closed^. (**d**) SEC traces of UBXD1 mutants alone or incubated with p97 and ADP, showing a left shift in peak elution volume for p97 samples with UBXD1. Fractions in the shaded range were analyzed by SDS-PAGE. (**e**) Coomassie Brilliant Blue-stained SDS-PAGE gels of fractions from SEC runs in (d). (**f**) Cryo-EM processing workflow for the p97/UBXD1^LX^/ADP dataset. (**g**) FSC curve and particle orientation distribution plot for p97:UBXD1^LX^. (**h**) Sharpened density map colored by local resolution (0.143 cutoff) of p97:UBXD1^LX^. (**i**) Map-model FSC for p97:UBXD1^LX^. Displayed resolution was determined using the masked map. (**j**) Cryo-EM processing workflow for the p97/UBXD1^H4X^/ADP dataset. (**k**) Unsharpened map of class 1 from p97/UBXD1^LX^/ADP dataset. (l) Unsharpened map of class 1 from p97/UBXD1^H4X^/ADP. (**m**) Top: unsharpened map of p97:UBXD1^LX^. Note lack of density for the D2 domain of the protomer clockwise from the VIM/H4 bound protomer. Bottom: filtered map, colored only by p97 density, confirming that density for the aforementioned D2 domain is present. (**n**) Top: sharpened map of p97:UBXD1^H4X^, colored as in Fig. 1. Bottom: filtered map, confirming that density for the PUB domain is present. (**o**) Positions of calculated centroids for D1 (residues 210-458, magenta spheres) and D2 domains (residues 483-763, green spheres) of P1 and P6, for p97^ADP^ and p97:UBXD1^open^.

**Extended Data Fig. 8.**
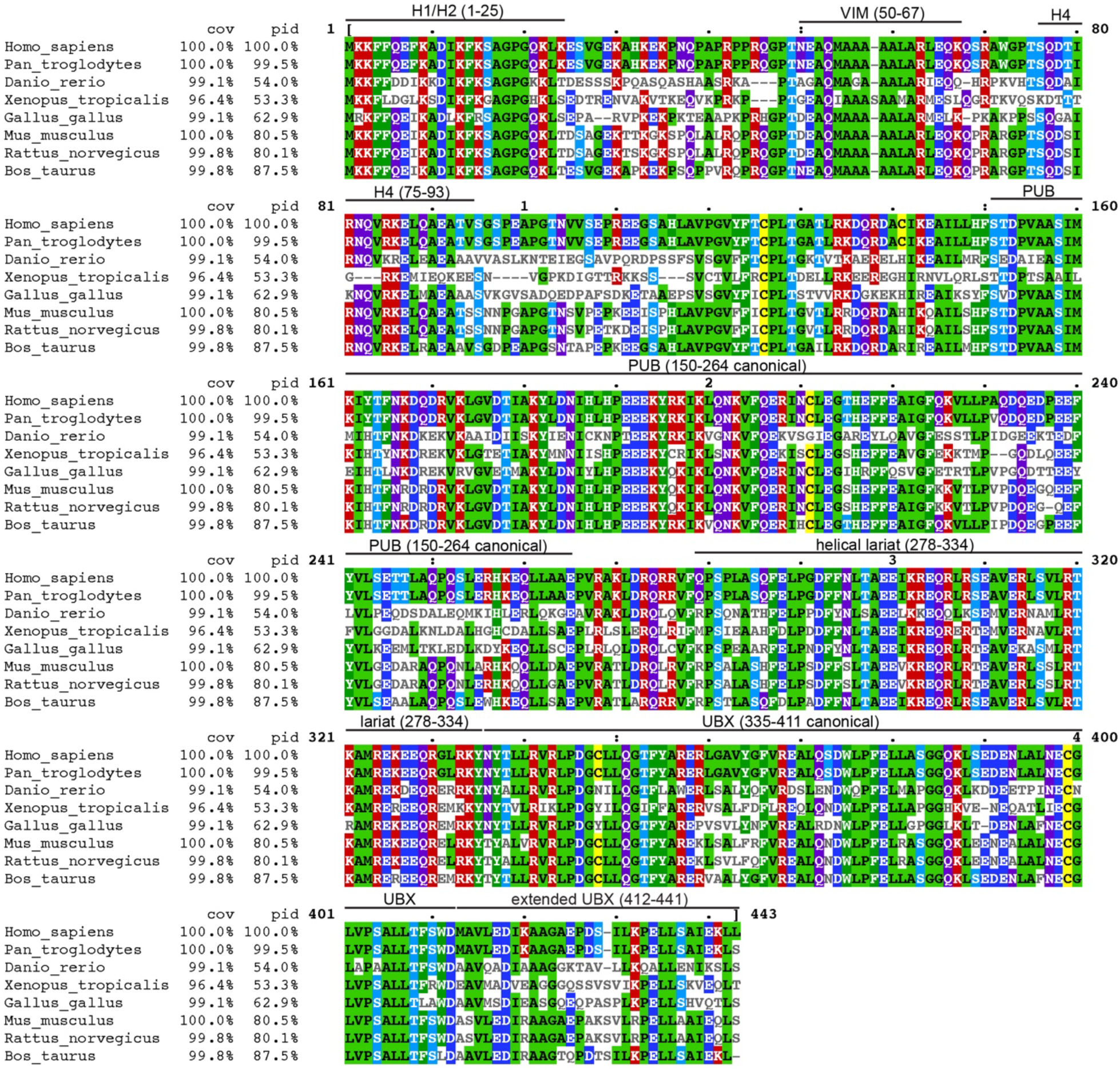
Alignment of UBXD1 sequences. Multiple sequence alignment of UBXD1 homologs from *Homo sapiens*, *Mus musculus*, *Xenopus tropicalis*, *Gallus gallus*, *Rattus norvegicus*, *Bos taurus*, *Pan troglodytes*, and *Danio rerio*. Structural elements and residue ranges (in the human sequence) are marked above. Cov = covariace relative to the human sequence, Pid = percent identity relative to the human sequence.

**Extended Data Table 1.** Cryo-EM data collection, refinement, and validation statistics of p97:UBXD1^LX^.

|  |  |
| --- | --- |
|  | p97:UBXD1<sup>LX</sup><br>(EMD-28992,<br>PDB 8FCT) |
| **Data collection and processing** |  |
| Microscope and camera | Titan Krios, K3 |
| Magnification | 59,952 |
| Voltage (kV) | 300 |
| Data acquisition software | SerialEM |
| Exposure navigation | Image shift |
| Electron exposure (e<sup>-</sup>/Å<sup>2</sup>) | 43 |
| Defocus range (μm) | -0.5 to -2.0 |
| Pixel size (Å) | 0.834 |
| Symmetry imposed | C1 |
| Initial particle images (no.) | 2,250,090 |
| Final particle images (no.) | 106,993 |
| Map resolution (Å) | 3.4 |
| FSC threshold | 0.143 |
| Map resolution range (Å) | 2.5-10 |
| **Refinement** |  |
| Model resolution (Å) | 4.0 |
| FSC threshold | 0.5 |
| Map sharpening *B* factor (Å<sup>2</sup>) | -107.0 |
| Model composition |  |
| Nonhydrogen atoms | 34,329 |
| Protein residues | 4,356 |
| Ligands | 12 |
| *B* factors (Å<sup>2</sup>) |  |
| Protein | 26.10 |
| Ligand | 13.34 |
| R. m. s. deviations |  |
| Bond lengths (Å) | 0.011 |
| Bond angles (°) | 1.040 |
| **Validation** |  |
| MolProbity score | 0.71 |
| Clashscore | 0.62 |
| Poor rotamers (%) | 0.05 |
| Ramachandran plot |  |
| Favored (%) | 98.52 |
| Allowed (%) | 1.34 |
| Disallowed (%) | 0.14 |

## SUPPLEMENTARY INFORMATION

**Supplementary Video 1. p97:UBXD1^closed^ and p97:UBXD1^open^ states**

Sharpened maps and models for p97:UBXD1^closed^ and p97:UBXD1^open^ are shown.

**Supplementary Video 2. Transition between p97^ADP^, p97:UBXD1^closed^ and p97:UBXD1^open^**

Morphs between models of p97^ADP^ (PDB 5FTK), p97:UBXD1^closed^ and p97:UBXD1^open^ are shown.

**Supplementary Video 3. 3D variability analysis of p97:UBXD1^closed^ and p97:UBXD1^open^**

Volumes calculated from the first component of a 3D Variability Analysis job using particles in the closed and open states are shown, colored as in Figure 1. This identifies a clear hexamer opening and upward and downward rotation of protomers P6 and P1, respectively.

**Supplementary Video 4. H4 binding disrupts p97 D2 inter-protomer contacts**

Morph between p97:UBXD1^closed^ and p97:UBXD1^H4^ is shown, showing rotation of bound D2 domain and disruption of contact with α5’ (red) of clockwise adjacent protomer.

**Supplementary Video 5. 3D variability analysis of p97:UBXD1^LX^**

Volumes calculated from the third principal component of a 3D Variability Analysis job using particles from class 2 of the p97:UBXD1^LX^ dataset are shown. This reveals a transition from a p97 hexamer without appreciable density corresponding to UBXD1 and ordered D2 domains to a hexamer with strong VIM and H4 density. In this state, the H4-bound D2 domain rotates upward and density for the D2 domain of the clockwise protomer disappears, indicating flexibility.

**Supplementary Video 6. Summary of p97:UBXD1 interactions**

UBXD1 domains are shown binding to the p97 hexamer in p97:UBXD1^H4^.

## REFERENCES

1. Stach, L. & Freemont, P. S. The AAA+ ATPase p97, a cellular multitool. Biochem J 474, 2953–2976 (2017).

2. Boom, J. van den & Meyer, H. VCP/p97-Mediated Unfolding as a Principle in Protein Homeostasis and Signaling. Mol Cell 69, 182–194 (2018).

3. Watts, G. D. J. et al. Inclusion body myopathy associated with Paget disease of bone and frontotemporal dementia is caused by mutant valosin-containing protein. Nat Genet 36, 377– 381 (2004).

4. Darwich, N. F. et al. Autosomal dominant VCP hypomorph mutation impairs disaggregation of PHF-tau. Science 370, (2020).

5. Johnson, J. O. et al. Exome Sequencing Reveals VCP Mutations as a Cause of Familial ALS. Neuron 68, 857–864 (2010).

6. Huryn, D. M., Kornfilt, D. J. P. & Wipf, P. p97: An Emerging Target for Cancer, Neurodegenerative Diseases, and Viral Infections. J Med Chem 63, 1892–1907 (2019).

7. Fessart, D., Marza, E., Taouji, S., Delom, F. & Chevet, E. P97/CDC-48: Proteostasis control in tumor cell biology. Cancer Lett 337, 26–34 (2013).

8. Cooney, I. et al. Structure of the Cdc48 segregase in the act of unfolding an authentic substrate. Science 365, 502–505 (2019).

9. Twomey, E. C. et al. Substrate processing by the Cdc48 ATPase complex is initiated by ubiquitin unfolding. Science 365, eaax1033 (2019).

10. Stein, A., Ruggiano, A., Carvalho, P. & Rapoport, T. A. Key Steps in ERAD of Luminal ER Proteins Reconstituted with Purified Components. Cell 158, 1375–1388 (2014).

11. Brandman, O. et al. A Ribosome-Bound Quality Control Complex Triggers Degradation of Nascent Peptides and Signals Translation Stress. Cell 151, 1042–1054 (2012).

12. Papadopoulos, C. et al. VCP/p97 cooperates with YOD1, UBXD1 and PLAA to drive clearance of ruptured lysosomes by autophagy. Embo J 36, 135–150 (2016).

13. Tanaka, A. et al. Proteasome and p97 mediate mitophagy and degradation of mitofusins induced by Parkin. J Cell Biology 191, 1367–1380 (2010).

14. Ritz, D. et al. Endolysosomal sorting of ubiquitylated caveolin-1 is regulated by VCP and UBXD1 and impaired by VCP disease mutations. Nat Cell Biol 13, 1116–1123 (2011).

15. Buchberger, A., Schindelin, H. & Hänzelmann, P. Control of p97 function by cofactor binding. Febs Lett 589, 2578–2589 (2015).

16. Hänzelmann, P. & Schindelin, H. The Interplay of Cofactor Interactions and Post-translational Modifications in the Regulation of the AAA+ ATPase p97. Frontiers Mol Biosci 4, 21 (2017).

17. Bento, A. C. et al. UBXD1 is a mitochondrial recruitment factor for p97/VCP and promotes mitophagy. Sci Rep-uk 8, 12415 (2018).

18. DeLaBarre, B. & Brunger, A. T. Complete structure of p97/valosin-containing protein reveals communication between nucleotide domains. Nat Struct Mol Biol 10, 856–863 (2003).

19. Banerjee, S. et al. 2.3 Å resolution cryo-EM structure of human p97 and mechanism of allosteric inhibition. Science 351, 871–875 (2016).

20. Huyton, T. et al. The crystal structure of murine p97/VCP at 3.6Å. J Struct Biol 144, 337–348 (2003).

21. Zhang, X. et al. Structure of the AAA ATPase p97. Mol Cell 6, 1473–1484 (2000).

22. Rouiller, I. et al. Conformational changes of the multifunction p97 AAA ATPase during its ATPase cycle. Nat Struct Biol 9, 950–957 (2002).

23. Tang, W. K. et al. A novel ATP-dependent conformation in p97 N–D1 fragment revealed by crystal structures of disease-related mutants. Embo J 29, 2217–2229 (2010).

24. DeLaBarre, B. & Brunger, A. T. Nucleotide Dependent Motion and Mechanism of Action of p97/VCP. J Mol Biol 347, 437–452 (2005).

25. Pan, M. et al. Mechanistic insight into substrate processing and allosteric inhibition of human p97. Nat Struct Mol Biol 28, 614–625 (2021).

26. Puchades, C. et al. Structure of the mitochondrial inner membrane AAA+ protease YME1 gives insight into substrate processing. Science 358, eaao0464 (2017).

27. Puchades, C., Sandate, C. R. & Lander, G. C. The molecular principles governing the activity and functional diversity of AAA+ proteins. Nat Rev Mol Cell Bio 21, 43–58 (2020).

28. Gates, S. N. et al. Ratchet-like polypeptide translocation mechanism of the AAA+ disaggregase Hsp104. Science 357, 273–279 (2017).

29. Rizo, A. N. et al. Structural basis for substrate gripping and translocation by the ClpB AAA+ disaggregase. Nat Commun 10, 2393 (2019).

30. Bodnar, N. O. & Rapoport, T. A. Molecular Mechanism of Substrate Processing by the Cdc48 ATPase Complex. Cell 169, 722–735.e9 (2017).

31. Bodnar, N. O. et al. Structure of the Cdc48 ATPase with its ubiquitin-binding cofactor Ufd1-Npl4. Nat Struct Mol Biol 25, 616–622 (2018).

32. Weith, M. et al. Ubiquitin-Independent Disassembly by a p97 AAA-ATPase Complex Drives PP1 Holoenzyme Formation. Mol Cell 72, 766–777.e6 (2018).

33. Kracht, M. et al. Protein Phosphatase-1 Complex Disassembly by p97 is Initiated through Multivalent Recognition of Catalytic and Regulatory Subunits by the p97 SEP-domain Adapters. J Mol Biol 432, 6061–6074 (2020).

34. Boom, J. van den et al. Targeted substrate loop insertion by VCP/p97 during PP1 complex disassembly. Nat Struct Mol Biol 28, 964–971 (2021).

35. Haines, D. S. et al. Protein Interaction Profiling of the p97 Adaptor UBXD1 Points to a Role for the Complex in Modulating ERGIC-53 Trafficking*. Mol Cell Proteomics 11, M111.016444 (2012).

36. Trusch, F. et al. The N-terminal Region of the Ubiquitin Regulatory X (UBX) Domain-containing Protein 1 (UBXD1) Modulates Interdomain Communication within the Valosin-containing Protein p97. J Biol Chem 290, 29414–27 (2015).

37. Schuetz, A. K. & Kay, L. E. A Dynamic molecular basis for malfunction in disease mutants of p97/VCP. Elife 5, e20143 (2016).

38. Huang, R., Ripstein, Z. A., Rubinstein, J. L. & Kay, L. E. Cooperative subunit dynamics modulate p97 function. Proc National Acad Sci 116, 201815495 (2018).

39. Kern, M., Fernandez-Sáiz, V., Schäfer, Z. & Buchberger, A. UBXD1 binds p97 through two independent binding sites. Biochem Bioph Res Co 380, 303–307 (2009).

40. Blueggel, M., Boom, J. van den, Meyer, H., Bayer, P. & Beuck, C. Structure of the PUB Domain from Ubiquitin Regulatory X Domain Protein 1 (UBXD1) and Its Interaction with the p97 AAA+ ATPase. Biomol 9, 876 (2019).

41. Hänzelmann, P. & Schindelin, H. The Structural and Functional Basis of the p97/Valosin-containing Protein (VCP)-interacting Motif (VIM) MUTUALLY EXCLUSIVE BINDING OF COFACTORS TO THE N-TERMINAL DOMAIN OF p97*. J Biol Chem 286, 38679–38690 (2011).

42. Stapf, C., Cartwright, E., Bycroft, M., Hofmann, K. & Buchberger, A. The General Definition of the p97/Valosin-containing Protein (VCP)-interacting Motif (VIM) Delineates a New Family of p97 Cofactors*. J Biol Chem 286, 38670–38678 (2011).

43. Madsen, L. et al. Ubxd1 is a novel co-factor of the human p97 ATPase. Int J Biochem Cell Biology 40, 2927–2942 (2008).

44. Jumper, J. et al. Highly accurate protein structure prediction with AlphaFold. Nature 596, 583–589 (2021).

45. Varadi, M. et al. AlphaFold Protein Structure Database: massively expanding the structural coverage of protein-sequence space with high-accuracy models. Nucleic Acids Res 50, D439– D444 (2021).

46. Arumughan, A. et al. Quantitative interaction mapping reveals an extended UBX domain in ASPL that disrupts functional p97 hexamers. Nat Commun 7, 13047 (2016).

47. Bulfer, S. L., Chou, T.-F. & Arkin, M. R. p97 Disease Mutations Modulate Nucleotide-Induced Conformation to Alter Protein–Protein Interactions. Acs Chem Biol 11, 2112–2116 (2016).

48. Blythe, E. E., Gates, S. N., Deshaies, R. J. & Martin, A. Multisystem Proteinopathy Mutations in VCP/p97 Increase NPLOC4·UFD1L Binding and Substrate Processing. Structure 27, 1820–1829.e4 (2019).

49. Shorter, J. & Southworth, D. R. Spiraling in Control: Structures and Mechanisms of the Hsp104 Disaggregase. Csh Perspect Biol 11, a034033 (2019).

50. Holm, L. Structural Bioinformatics, Methods and Protocols. Methods Mol Biology 2112, 29–42 (2020).

51. Orme, C. M. & Bogan, J. S. The Ubiquitin Regulatory X (UBX) Domain-containing Protein TUG Regulates the p97 ATPase and Resides at the Endoplasmic Reticulum-Golgi Intermediate Compartment*. J Biol Chem 287, 6679–6692 (2012).

52. Nagahama, M. et al. SVIP Is a Novel VCP/p97-interacting Protein Whose Expression Causes Cell Vacuolation. Mol Biol Cell 14, 262–273 (2003).

53. Ballar, P. et al. Identification of SVIP as an Endogenous Inhibitor of Endoplasmic Reticulum-associated Degradation. J Biol Chem 282, 33908–33914 (2007).

54. Johnson, A. E. et al. SVIP is a molecular determinant of lysosomal dynamic stability, neurodegeneration and lifespan. Nat Commun 12, 513 (2021).

55. Chou, T.-F. et al. Specific Inhibition of p97/VCP ATPase and Kinetic Analysis Demonstrate Interaction between D1 and D2 ATPase Domains. J Mol Biol 426, 2886–2899 (2014).

56. Schütz, A. K., Rennella, E. & Kay, L. E. Exploiting conformational plasticity in the AAA+ protein VCP/p97 to modify function. Proc National Acad Sci 114, E6822–E6829 (2017).

57. Wang, Q., Song, C., Yang, X. & Li, C.-C. H. D1 Ring Is Stable and Nucleotide-independent, whereas D2 Ring Undergoes Major Conformational Changes during the ATPase Cycle of p97-VCP*. J Biol Chem 278, 32784–32793 (2003).

58. Petrović, S., et al. Structural remodeling of AAA+ ATPase p97 by adaptor protein ASPL facilitates posttranslational methylation by METTL21D. Proc National Acad Sci 120, e2208941120 (2023).

59. Buchberger, A., Howard, M. J., Proctor, M. & Bycroft, M. The UBX domain: a widespread ubiquitin-like module11Edited by P. E. Wright. J Mol Biol 307, 17–24 (2001).

60. Blythe, E. E., Olson, K. C., Chau, V. & Deshaies, R. J. Ubiquitin- and ATP-dependent unfoldase activity of P97/VCP•NPLOC4•UFD1L is enhanced by a mutation that causes multisystem proteinopathy. Proc National Acad Sci 114, E4380–E4388 (2017).

61. Mastronarde, D. N. Automated electron microscope tomography using robust prediction of specimen movements. J Struct Biol 152, 36–51 (2005).

62. Zheng, S. Q. et al. MotionCor2: anisotropic correction of beam-induced motion for improved cryo-electron microscopy. Nat Methods 14, 331–332 (2017).

63. Punjani, A., Rubinstein, J. L., Fleet, D. J. & Brubaker, M. A. cryoSPARC: algorithms for rapid unsupervised cryo-EM structure determination. Nat Methods 14, 290–296 (2017).

64. Scheres, S. H. W. RELION: Implementation of a Bayesian approach to cryo-EM structure determination. J Struct Biol 180, 519–530 (2012).

65. Pettersen, E. F. et al. UCSF Chimera—A visualization system for exploratory research and analysis. J Comput Chem 25, 1605–1612 (2004).

66. Croll, T. I. ISOLDE: a physically realistic environment for model building into low-resolution electron-density maps. Acta Crystallogr Sect D Struct Biology 74, 519–530 (2018).

67. Pettersen, E. F. et al. UCSF ChimeraX: Structure visualization for researchers, educators, and developers. Protein Sci 30, 70–82 (2021).

68. Emsley, P., Lohkamp, B., Scott, W. G. & Cowtan, K. Features and development of Coot. Acta Crystallogr Sect D Biological Crystallogr 66, 486–501 (2010).

69. Afonine, P. V. et al. Real-space refinement in PHENIX for cryo-EM and crystallography. Acta Crystallogr Sect D Struct Biology 74, 531–544 (2018).

70. Madeira, F. et al. Search and sequence analysis tools services from EMBL-EBI in 2022. Nucleic Acids Res 50, W276–W279 (2022).

